# Extracellular matrix assembly stress drives *Drosophila* central nervous system morphogenesis

**DOI:** 10.1101/2022.04.18.488510

**Authors:** Eduardo Serna-Morales, Besaiz J. Sánchez-Sánchez, Stefania Marcotti, Anushka Bhargava, Anca Dragu, Liisa M. Hirvonen, María-del-Carmen Díaz-de-la-Loza, Matyas Mink, Susan Cox, Emily Rayfield, Brian M. Stramer

**Affiliations:** Randall Centre for Cell and Molecular Biophysics, King’s College London; London, UK; Institute of Medical Biology, University of Szeged, Szeged, Hungary; School of Earth Sciences, University of Bristol, Bristol, UK

## Abstract

The forces controlling tissue morphogenesis are attributed to cellular-driven activities and any role for extracellular matrix (ECM) is assumed to be passive. However, all polymer networks, including ECM, can theoretically develop autonomous stresses during their assembly. Here we examine the morphogenetic function of an ECM prior to reaching homeostatic equilibrium by analyzing *de novo* ECM assembly during *Drosophila* ventral nerve cord (VNC) condensation. Asymmetric VNC shortening and a rapid decrease in surface area correlate with exponential assembly of Collagen-IV (Col4) surrounding the tissue. Concomitantly, a transient developmentally-induced Col4 gradient leads to coherent long-range flow of ECM, which equilibrates the Col4 network. Finite element analysis and perturbation of Col4 network formation through the generation of dominant Col4-truncations that affect assembly, reveals that VNC morphodynamics is driven by a sudden increase in ECM-driven surface tension. These data highlight that ECM assembly stress and associated network instabilities can actively participate in tissue morphogenesis.

## Introduction

The forces controlling tissue development are attributed to cellular-driven activities (Gilbert, 2000). Any role for ECM in modulating overall tissue shape is largely assumed to be passive by providing a substrate that allows cells to actively remodel their environment (Walma and Yamada, 2020). However, recent work on developing semicircular canals in zebrafish has revealed that ECM accumulation can have a more instructive role in shaping tissues through swelling and modulation of osmotic pressure (Munjal et al., 2021). Additionally, all polymer networks – including ECM – can theoretically develop autonomous stresses during their initial polymerisation, and there has been speculation that the out of equilibrium ECM behaviours may be playing a more active role during tissue morphogenesis than many would assume (Loganathan et al., 2016; Newman et al., 1987; Newman et al., 1985; Zamir et al., 2008). Investigating these active mechanisms requires analyzing the ECM network during embryonic assembly prior to reaching a crosslinked, homeostatic state. Here we exploit *Drosophila* embryogenesis, which involves a *de novo* burst of ECM polymerization midway through development (Matsubayashi et al., 2017; Matsubayashi et al., 2020), to examine the role of an out of equilibrium ECM network during central nervous system morphogenesis.

## Results

### Drosophila VNC condensation consists of two distinct temporal phases, and initial anisotropic changes in tissue morphology are independent of VNC cellular activity

During stage 14 of *Drosophila* development the embryonic nerve cord begins to undergo a sudden reduction in length in a process called VNC condensation. Time-lapse imaging of the entire ~12 hours of condensation revealed two distinct phases in VNC morphodynamics: a rapid 1^st^ phase when most of the morphological remodeling occurs in which the tissue shortens asymmetrically from tail to head over ~3 hours, and a slower 2^nd^ phase resulting in a symmetric reduction in length over the remaining ~9 hours of development (Figure 1A and S1A,B; Movie 1-part 1). Analysis of tissue geometry revealed that while the 2^nd^ phase involved a reduction in tissue volume, the change in shape during the 1^st^ phase was isovolumetric suggesting that, at least for the 1^st^ phase, the process is not a simple ‘condensation’ (Figure 1B,C). Interestingly, it was recently reported that the scaling of VNC length with embryo length is suddenly lost after the start of condensation (Tiwari et al., 2021), which is likely explained by the increase in thickness of the tissue (Figure 1B,C). These data suggest that VNC morphogenesis involves temporally controlled and mechanistically distinct processes.

**Fig. 1.**
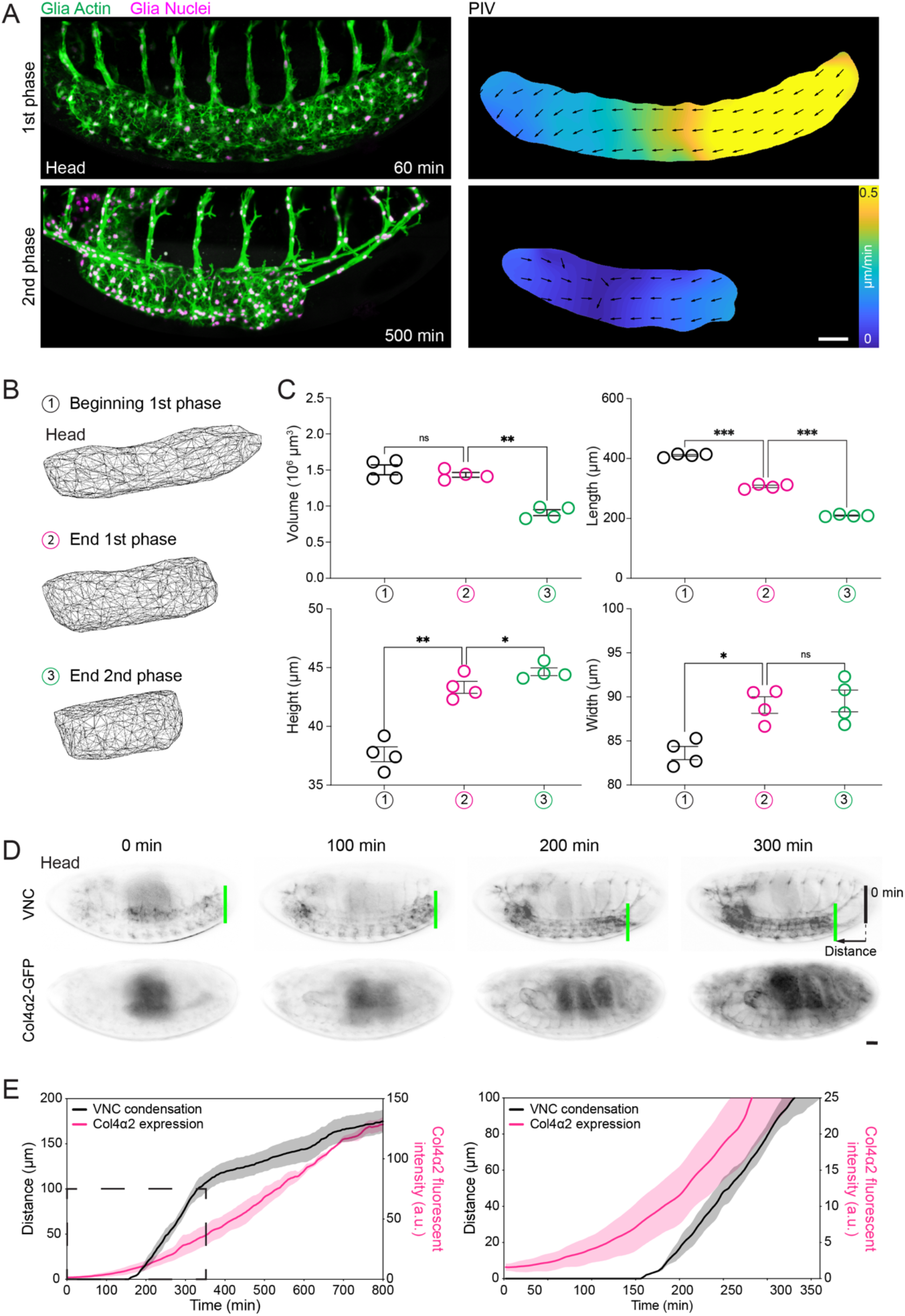
VNC morphogenesis involves distinct stages and correlates with the initiation of Col4 assembly. (**A**) Live imaging of VNC morphogenesis (left panels) and quantification of tissue motion by PIV (right panels) revealing anisotropic (1^st^ phase) and isotropic (2^nd^ phase) phases of condensation. (**B**) 3D reconstruction of VNC shape during the different phases of condensation. (**C**) Quantification of VNC shape at the start (1), end of the 1^st^ phase (2) and end of the 2^nd^ phase (3) of condensation. Repeated measures one-way ANOVA with Geisser-Greenhouse correction and Holm-Šídák’s multiple comparisons test. N = 4 embryos. (**D**) Live imaging of VNC condensation (highlighted by the green and black lines) and the induction of Col4 production by quantifying fluorescence intensity of Col4α2-GFP. (**E**) Correlation of Col4 fluorescence intensity with the rate of VNC condensation as measured by tracking the motion of the tail of the tissue (i.e., the difference between the green and black lines in (D)). Right panel focuses on the data highlighted by the dashed square. Scale bars = 30 μm, N = 3 embryos.

The VNC is composed of a central nerve bundle surrounded by supporting glial cells, and previous work revealed that affecting glial or neuronal activity perturbs VNC condensation (Olofsson and Page, 2005). However, inhibiting cell activity by driving a dominant negative (DN) Myosin II or Rac GTPase in glia or neurons led to relatively minor effects on condensation whereby the 1^st^ phase was left intact (Figure S1C-H). Additionally, tracking glial movement during the 1^st^ phase revealed a coherent flow of cells without any cellular rearrangements that could explain the VNC morphodynamics (Figure S1I-L; Movie 1-part 2). These data suggest that there are VNC cell-extrinsic forces driving the 1^st^ phase of condensation and we therefore set out to determine what was initiating the process.

### Initiation of VNC condensation is correlated with hemocyte deposition of Col4 on the tissue surface and a sudden increase in tissue stiffness

There is an underexplored function of the ECM during *Drosophila* VNC morphogenesis. The VNC is ensheathed by a basement membrane (BM) composed of Collagen Type IV (Col4) and Laminin, and mutations in BM components perturb condensation (Borchiellini et al., 1996; Urbano et al., 2009). Correlation of VNC condensation with Col4 levels using a GFP-trap in the Col4α2 subunit revealed that initiation of the process coincided with embryonic induction of Col4 expression and an exponential increase in production (Figure 1D,E; Movies 1-parts3,4). In contrast, Laminin is produced at much earlier stages of embryogenesis (Matsubayashi *et al*., 2017; Matsubayashi *et al*., 2020), which led us to hypothesize that Col4 assembly was specifically involved in triggering the start of VNC morphogenesis.

*Drosophila* hemocytes (macrophages), which are also hypothesized to be involved in VNC condensation (Matsubayashi *et al*., 2017; Olofsson and Page, 2005), are the major producers of Col4 during embryogenesis (Matsubayashi *et al*., 2017) and their developmental dispersal from their birth in the head mesoderm coincides with the start of condensation (Movie 1-part 5). Hemocytes deposit Col4 around the VNC as they disperse evenly throughout the embryo (Matsubayashi *et al*., 2017), which consequently results in a transient gradient of Col4 from head to tail along the tissue (Figure S2A,B). Subsequently, as condensation proceeds, the Col4 gradient equilibrated such that by the start of the 2^nd^ phase there was an even distribution of Col4 around the VNC (Figure S2A,B). This induction in Col4 also correlated with an increase in VNC stiffness from the 1^st^ to the 2^nd^ phase of condensation suggesting that ECM assembly was suddenly altering the mechanical properties of the tissue (Figure 2A).

**Fig. 2.**
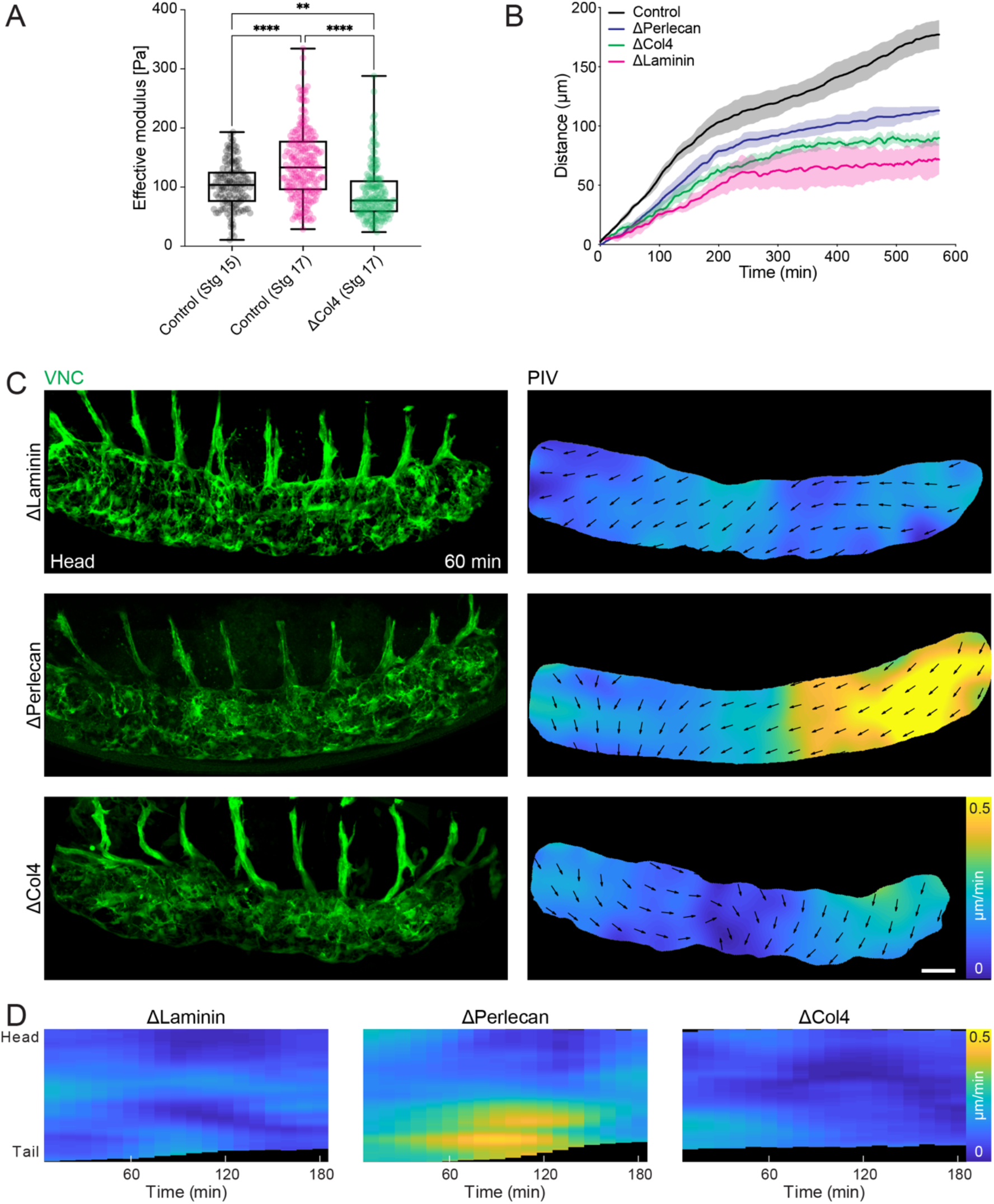
Col4 induction correlates with an increase in VNC stiffness and loss of Col4 inhibits the anisotropic phase of VNC condensation. (**A**) Atomic force microscopy of the VNC revealing an increase in stiffness from stage 15 to 17, which is lost in the absence of Col4. Kruskal-Wallis test and Dunn’s multiple comparisons test. N = 137 indentations over one Control embryo (stg 15), N = 207 indentations over two Control (stg 17) embryos, and N = 181 indentations over two DCol4 (stg 17) embryos. (**B**) Quantification of the rate of VNC condensation by tracking tail motion as in Fig. 1D,E in control (N = 4 embryos) and *laminin, perlecan*, and *col4* mutant embryos (N = 3 embryos for each mutation). (**C**) Live imaging of the 1^st^ phase of VNC condensation as in Fig. 1A in *laminin, perlecan*, and *col4* mutants. Scale bar = 30 μm. (**D**) Kymograph of the average speed of VNC condensation from PIV analysis in (C) highlighting an absence of an anisotropic phase of condensation in *laminin* and *col4* mutants.

### Inhibiting Col4 deposition or altering its distribution along the VNC surface prevents the initial anisotropic change in VNC morphology

Analysis of BM mutants revealed that in the absence of Laminin, which is required for subsequent BM component assembly (Matsubayashi *et al*., 2017), condensation is severely affected and the VNC falls apart due to cell clumping (Figure 2B-D; Movie 2-part 1). In contrast, in the absence of Perlecan, which is expressed slightly later during embryogenesis than Col4 (Matsubayashi *et al*., 2017), condensation was slower and the 2^nd^ phase severely affected, yet there was a clear asymmetric 1^st^ phase of VNC morphogenesis (Figure 2B-D; Movie 2-part 1). However, Col4 mutants had a distinct phenotype: Col4 mutant VNCs remained intact, yet deformed, and both phases of VNC condensation were severely inhibited (Figure 2B-D; Movie 2-part 1). Importantly, the 1^st^ phase of condensation was lost in the absence of Col4 and what little condensation was present was isotropic rather than asymmetric (Figure 2C,D; Movie 2-part 1). Additionally, Col4 mutants failed to show an increase in tissue stiffness (Figure 2A). The Col4 mutant VNC phenotype was specific to Col4 assembly around the tissue as local disruption of Col4 by expression of a surface-bound matrix metalloprotease (*Drosophila* MMP2) (Pastor-Pareja et al., 2008; Sui et al., 2018) with the same glial-Gal4 driver used to inhibit glial cell activity, phenocopied the Col4 mutant (Figure S2C,D). Additionally, preventing hemocyte release from the head of the embryo by expression of Rac DN, which causes a relative overabundance of Col4 towards the head (Matsubayashi *et al*., 2017), or genetically deleting hemocytes severely inhibited VNC shortening and deformed the tissue (Figure S3; Movies 2-parts 2,3). Therefore, similar to what was previously observed (Olofsson and Page, 2005; Page and Olofsson, 2008), disrupting hemocytes and Col4 distribution leads to a far more severe effect on VNC condensation than inhibiting VNC cellular activity.

### Finite element analysis reveals that the initial anisotropic change in VNC morphology can be explained by a sudden increase in asymmetric surface tension

Polymer networks, especially in thin film organization, are known to generate stresses during assembly, which can lead to polymer motion as stresses equilibrate (Freund and Suresh, 2009). Indeed, inhomogeneities in mixtures of ECM components in a test tube can generate forces that directly drive long-range translocation of inert particles (Newman *et al*., 1987; Newman *et al*., 1985). To determine whether BM-derived surface stress could contribute to the observed changes in VNC shape upon initiation of condensation we developed a finite element model consisting of two materials: an internal core representing the VNC attached to the brain, which is surrounded by a thin BM shell. Simulations revealed that a uniform pressure perpendicular to the surface was sufficient to decrease the length of the tissue from tail to head. However, this also led to constriction of the diameter of the VNC, which was not observed experimentally during the 1^st^ phase of condensation (Figure 3A, Appendix). In a second approach, we modeled the shell such that it instead exerted a surface tension, which can be modulated by altering its capacity to shrink and its stiffness. Consistent with the Col4 dependent increase in tissue stiffness during condensation, a simulated increase in surface tension asymmetrically reduced the tissue length, yet this also led to a slight reduction in tissue width, which was also not observed in real specimens (Figures 1B,C and 3A, Appendix). However, further simulations suggested that an anisotropic surface tension, which is predominantly along the length of the tissue, could explain the increase in width and height of the VNC during the 1^st^ phase of condensation (Figures 1B,C and 3A, Appendix). These data suggest that a BM assembly-dependent increase in tissue surface tension predominantly along the axis of the observed Col4 gradient can explain the rapid anisotropic change in tissue shape during initiation of VNC morphogenesis.

**Fig. 3.**
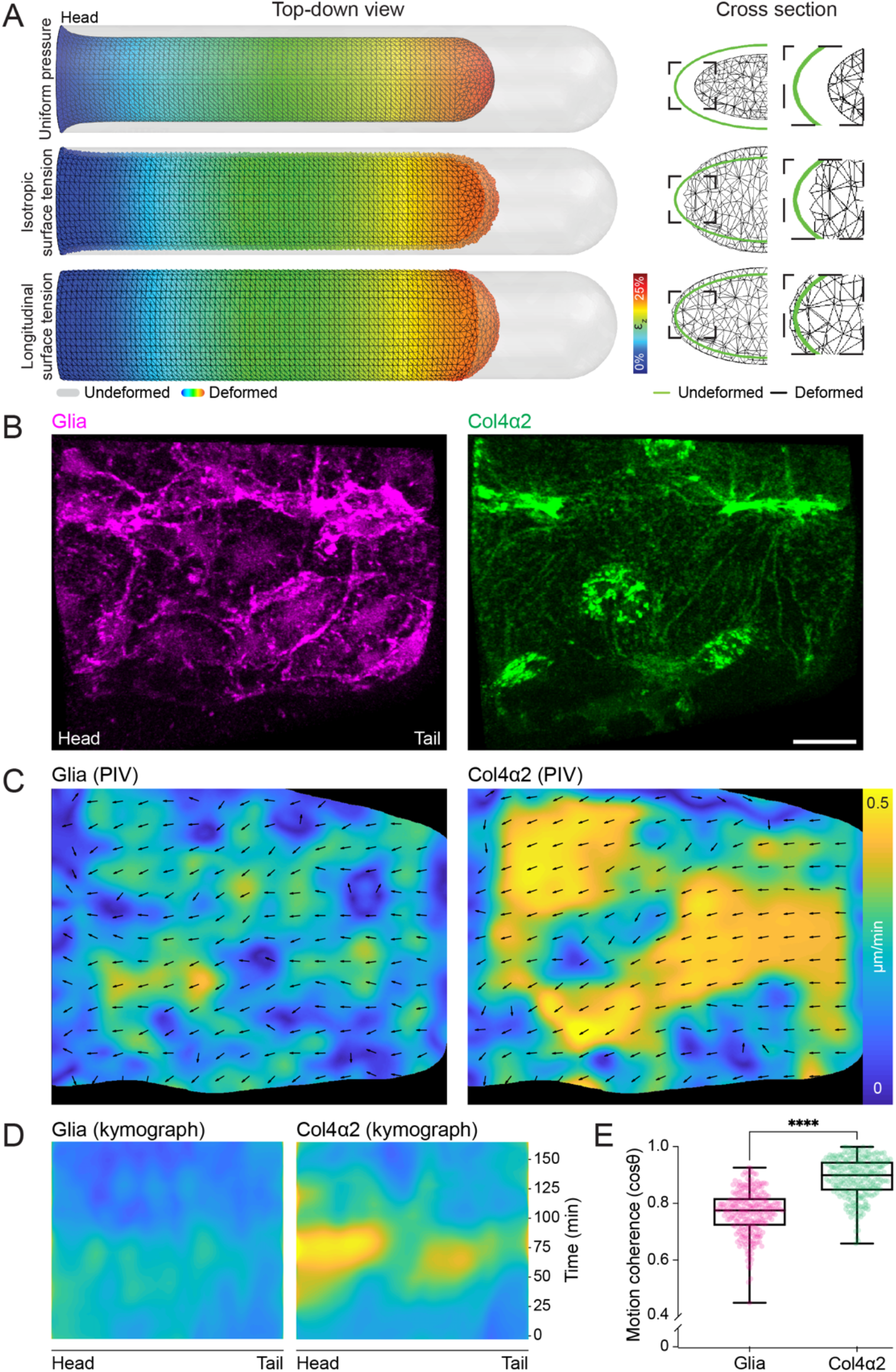
Anisotropy in surface tension and coherent long-range flow of Col4 is sufficient to explain the sudden isovolumetric change in VNC shape. (**A**) Finite element analysis of VNC morphogenesis. Simulations of VNC deformation assuming (top panel) a uniform normal surface pressure, (middle panel) a uniform increase in surface tension, or (bottom panel) an anisotropic increase in surface tension along the length of the tissue. Note that only the anisotropic surface tension leads to a reduction in length and an increase in height and width of the tissue. (**B**) Live imaging of glia and Col4 motion on the surface of the VNC. Scale bar = 10 μm. (**C**) Simultaneous tracking of glia and Col4 motion by PIV. (**D**) Kymograph of the PIV analysis in panel (C) highlighting a sudden increase in Col4 speed during VNC condensation which is not observed by tracking glia. (**E**) Correlation of the local alignment of PIV vectors reveals that the motion of the Col4 network is more coherent than glial motion. Mann-Whitney test. N = 256 vectors for each sample.

### Live imaging of Col4 assembly on the VNC surface reveals a coherent viscous-like flow along the axis of the predicted increase in tissue surface tension

We subsequently live imaged surface glial dynamics and Col4 accumulation surrounding the VNC at high spatiotemporal resolution to determine the possible interplay between cell activity and ECM remodelling. As VNC condensation initiated, a rapid and coherent long-range flow of Col4 was observed antiparallel to the Col4 gradient (*i.e*., tail to head direction), which was along the predicted predominant axis of surface tension (Figures 3B-D and S4A-C; Movie 3-parts 1,2). Simultaneously, there was a drift in the surface glia, which are in direct contact with the overlying ECM enveloping the tissue (Figure 3B,C; Movie 3-part 1). At the start of condensation, the surface glia are initially spaced out on the VNC surface and not in contact with each other, and eventually undergo a mesenchymal to epithelial transition to form an epithelial monolayer by the end of the 1^st^ phase (Schwabe et al., 2005; Schwabe et al., 2017). Despite the approximate 16% decrease in VNC surface area during the 1^st^ phase (determined from our measured change in tissue dimensions, Figure 1B,C) surface glial cells paradoxically increase in area as they spread themselves around the tissue (Schwabe *et al*., 2005; Schwabe *et al*., 2017), highlighting that glial cell contraction is not involved in this change in tissue geometry. We observed no obvious cell migration or cell rearrangements of glial cells that would explain the overall change in VNC morphology. Nor did we observe any local ECM remodelling by the underlying glial cells (Movie 3-part 1). Indeed, correlating the local alignment of particle image velocimetry (PIV) vectors of ECM and glial motion revealed that the Col4 network was moving more coherently over the tissue surface than the cells themselves (Figure 3D, E). Additionally, the Col4 network appeared to self-assemble with polymers stretching tens of microns linking the dispersed population of glial cells (Figure 3B). These data suggest that ECM-autonomous dynamics are involved in BM formation, which leads to long-range viscous flow and equilibration of the ECM network.

### Local disruption of the BM network or subtle perturbation of Col4 assembly inhibits the rate of VNC condensation

We next examined perturbations that affect ECM polymerization or ECM network organization on VNC morphodynamics. Simulations further suggested that local changes in surface tension should lead to global effects on the rate of condensation (Figure 4A, Appendix). To confirm the presence of long-range network effects, we locally cleaved the ECM network in a narrow stripe along the VNC using a parasegmental driver (Bowman et al., 2014) with MMP2 expression (Pastor-Pareja *et al*., 2008; Sui *et al*., 2018) and examined motion of the VNC by tracking the tail of the tissue. Cleaving the ECM network in a stripe in the middle of the VNC slowed the motion of the tail of the tissue, which was over a hundred microns distant to the site of cleavage (Figure 4B). This long-range effect suggests that a coherent and interconnected ECM network is required for VNC morphogenesis. We subsequently examined a temperature sensitive (TS) point mutant Col4 allele (G552D) (Kelemen-Valkony et al., 2012; Kiss et al., 2016) that resulted in the ECM network disintegrating when shifted to 29°C (Figure 4C,D). This revealed that subtle alteration of the temperature in this TS mutant was sufficient to affect the overall rate of VNC motion (Figure 4E). Additionally, driving a G552D Col4 transgene specifically in hemocytes severely inhibited VNC condensation confirming that hemocyte-derived Col4 is indeed essential for initiating the process (Figure S4D-F). We also generated an N-terminal truncation in the Col4α1 protein that removed part of the putative 7S domain (Figure 4F,G), which is hypothesised to be involved in multivalent interactions important for Col4 network assembly (Duncan et al., 1983; Siebold et al., 1987). Driving this transgene specifically in hemocytes was sufficient to perturb Col4 network formation and subtly affect the overall rate of condensation, which we speculate is due to a retardation of Col4 polymerization (Figure 4H,I).

**Fig. 4.**
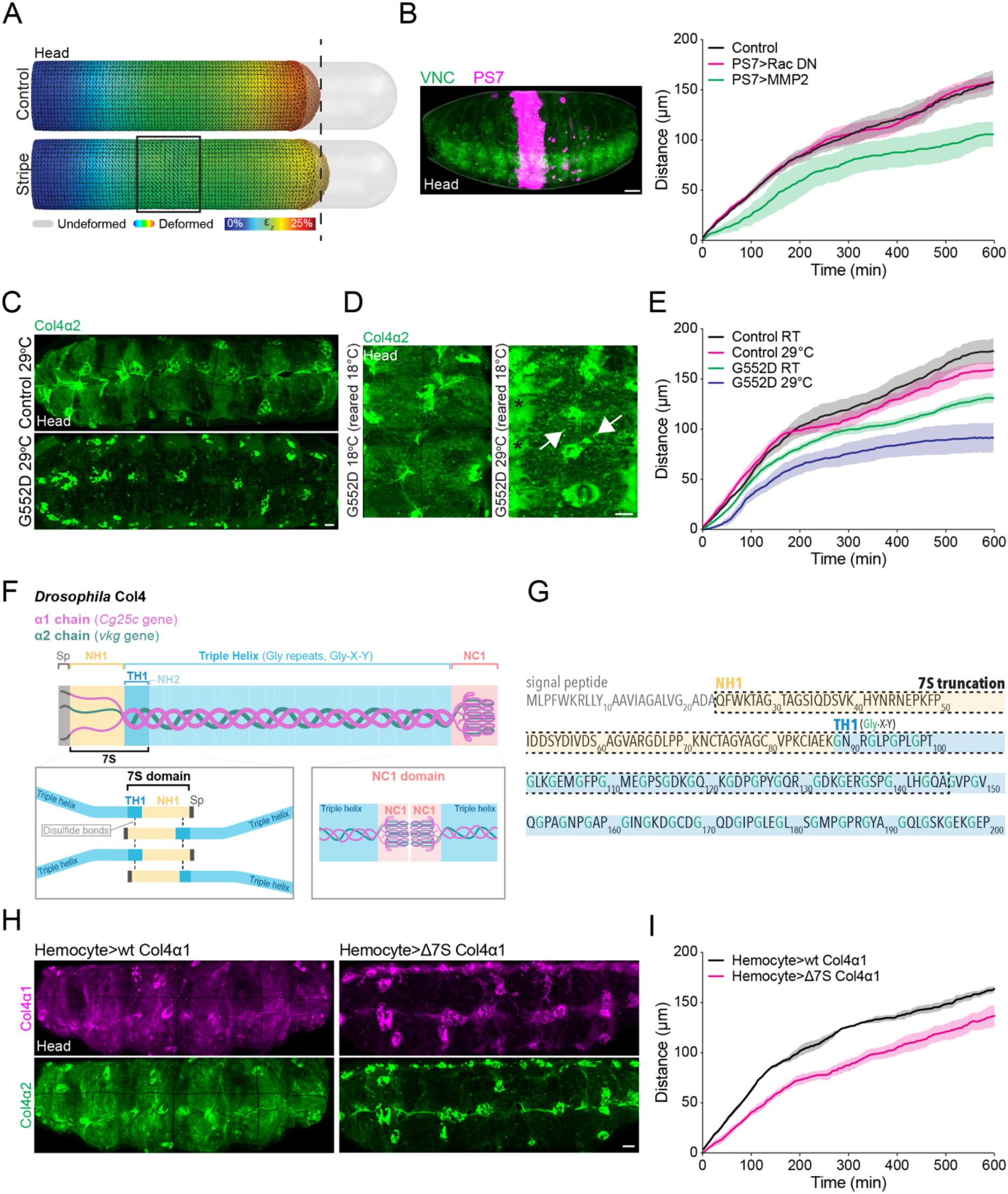
Direct perturbations of Col4 assembly slow the rate of VNC condensation. (**A**) Finite element analysis of VNC condensation in which surface tension is locally reduced in a stripe in the middle of the tissue. Simulations suggest that local perturbation of surface tension leads to a long-range reduction in deformation of the tail of the tissue. (**B**) Local disruption of the ECM network by expression of MMP2 in a central parasegment (PS7, N = 4 embryos), in contrast to Control (N = 3 embryos) and expression of Rac DN (N = 3 embryos), is sufficient to affect the rate of VNC condensation as measured by tracking the tail of the tissue as in Fig. 1D,E. Scale bar = 30 μm. (**C**) Live imaging of Col4α2-GFP reveals that rearing embryos containing a temperature sensitive (TS) point mutation in Col4α1 (G552D) at the non-permissive temperature (29°C) inhibits assembly of the BM network. Scale bar = 10 μm. (**D**) Rearing the TS mutant at the permissive temperature (18°C) to allow some ECM assembly and switching to the non-permissive temperature (29°C) leads to aggregation of extracellular Col4 (arrows) and accumulation of soluble Col4 in the hemocoel (asterisks) showing that the TS mutant affects the Col4 network. Scale bar = 10 μm. (**E**) Quantification of VNC condensation in the TS mutant by tracking the motion of the tail of the tissue reveals that the rate of condensation can be affected by temperature titration (RT: room temperature). Control data reused from Fig. 2B. N = 4 embryos for Control RT and Control 29°C, N = 5 embryos for G552D RT and G552D 29°C. **(F)** Schematic highlighting Col4 trimer domains and interactions hypothesized to drive network assembly. In the C-terminus of Col4 are interactions between NC1 domains, which lead to dimerization of Col4 trimers. At the N-terminus are interactions of a putative 7S domain, which consists of the first non-helical (NH1) and triple-helical (TH1) regions of the protein. Disulfide bonds and non-covalent interactions between 7S domains are hypothesized to lead to tetramerization of Col4 trimers. Sp, signal peptide. **(G)** Amino acid sequence of the N-terminus of *Drosophila* Col4. The dashed line highlights the amino acid sequence truncated from the putative 7S domain of Col4α1 to generate a dominant negative transgene. (**H**) Expression of wild-type (wt) Col4α1 or a truncation of the 7S domain in hemocytes shows that the deletion of the 7S domain leads to reduced incorporation into the ECM network and also reduces incorporation of Col4α2. Scale bar = 10 μm. (**I**) Quantification of VNC condensation after expression of the transgenes in (H) reveals that removal of the 7S domain slows the rate of motion of the tail of the tissue. N = 4 embryos for each sample.

## Discussion

The textbook view of morphogenesis has assumed that changes in tissue shape are driven by a limited repertoire cell-autonomous activities, such as cell rearrangements, shape changes, migrations, and divisions (Gilbert, 2000). However, the supramolecular nature of ECM networks, such as BMs, which rely on underexplored covalent and noncovalent interactions for their self-assembly (Horne-Badovinac, 2020; Keeley et al., 2020; Matsubayashi *et al*., 2020; Yurchenco and Furthmayr, 1984; Yurchenco et al., 1986), should not be neglected as sources of stress that may shape an embryo. Additionally, surface tension has been predicted to be a dominant force in shaping developing tissues (Thompson, 1961), and while alterations in surface tension are often assumed to be driven by epithelial activity (Lecuit and Lenne, 2007), BM networks are also likely to be involved. Indeed, in the case of VNC condensation, the sudden exponential increase in Col4 formation around the VNC is behaving as a compression sleeve, which actively envelops the tissue to shrink its surface area, resulting in an initial isovolumetric change in shape as surface stresses equilibrate. In contrast, in the 2^nd^ phase of condensation, once the asymmetric Col4 network stresses have dissipated, the cells within the VNC likely take a more active part in the process with the ECM possibly playing a more stereotypical passive role.

There are many reasons why a polymer network like the BM could generate forces during development. The polymerization process on its own, especially if there are instabilities such as concentration gradients during assembly, will lead to stress development and ECM-autonomous motion. Indeed, *in vitro* polymerization assays have revealed that a nonuniform distribution of collagen self-assembly can generate a surface tension gradient that leads to a Marangoni flow within the network (Nerger et al., 2020). Additionally, a mechanical mismatch between a polymer thin film, such as the BM, and the underlying substrate can also lead to material deformations (Francis et al., 2002; Freund and Suresh, 2009). As long-range coherent motion of ECM has been observed during a number of developmental processes (Loganathan *et al*., 2016; Szabo et al., 2011; Zamir *et al*., 2008), nonequilibrium phenomena in an immature ECM network in flux may be playing unappreciated active roles in tissue morphogenesis.

## Acknowledgments

We would like to thank Roberto Mayor, Nir Gov and Patrick Mesquida for comments and advice on the manuscript.

## Author contributions

Conceptualization: ESM, BJSS, SM, BMS

Methodology: ESM, BJSS, SM, MM, BMS

Investigation: ESM, BJSS, SM, AB, AD, LH, MD, MM, SC, ER, BMS

Funding acquisition: BMS

Project administration: ESM, BMS

Writing – original draft: BMS

Writing – review & editing: ESM, BJSS, SM, BMS

## Data and materials availability

All data are available in the main text or the supplementary materials.

## Supplementary Materials

Materials and Methods

Appendix

Figures and figure legends S1 to S4

Movie legends S1 to S3

Movies S1 to S3

## Materials and Methods

### Resource table

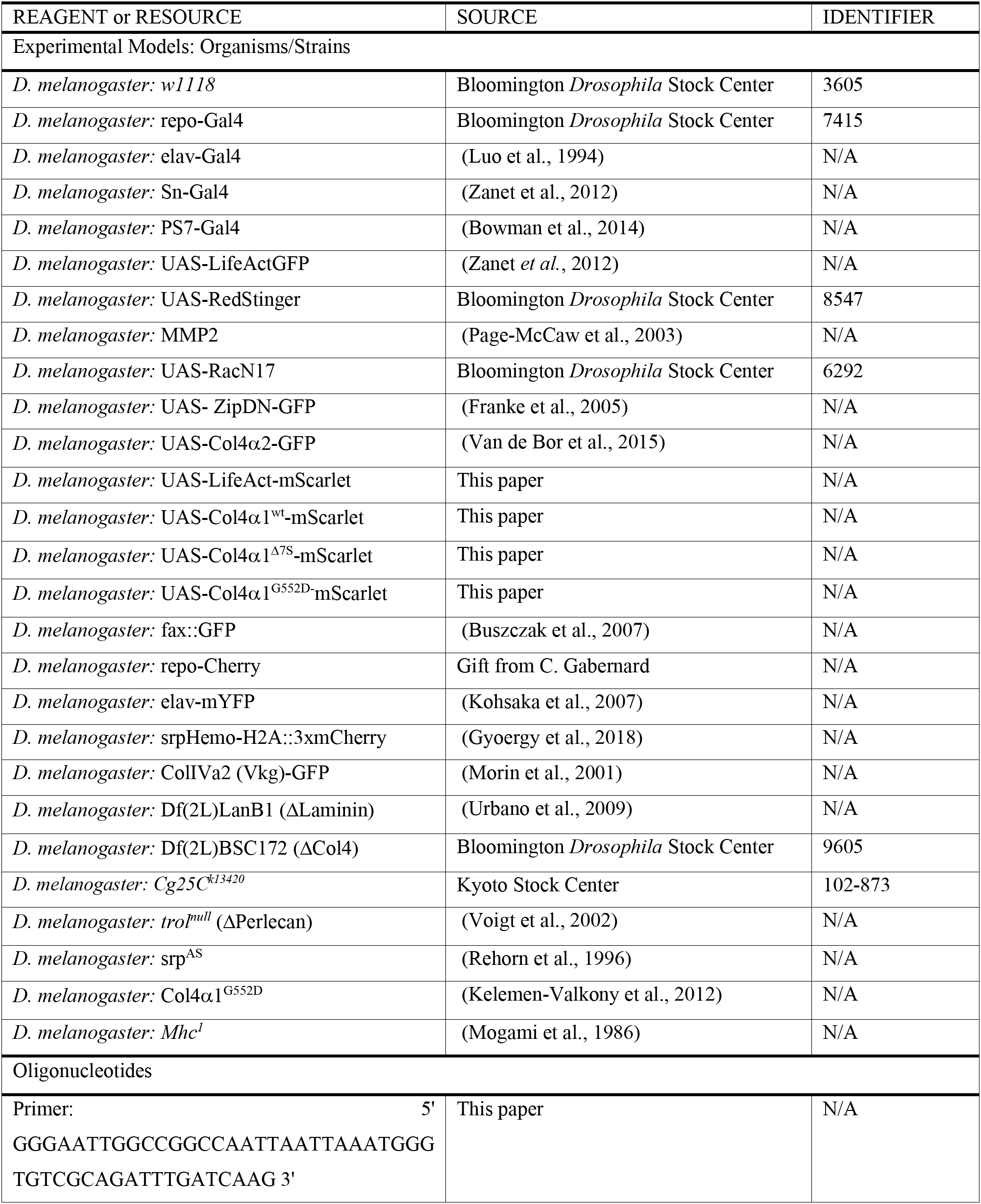
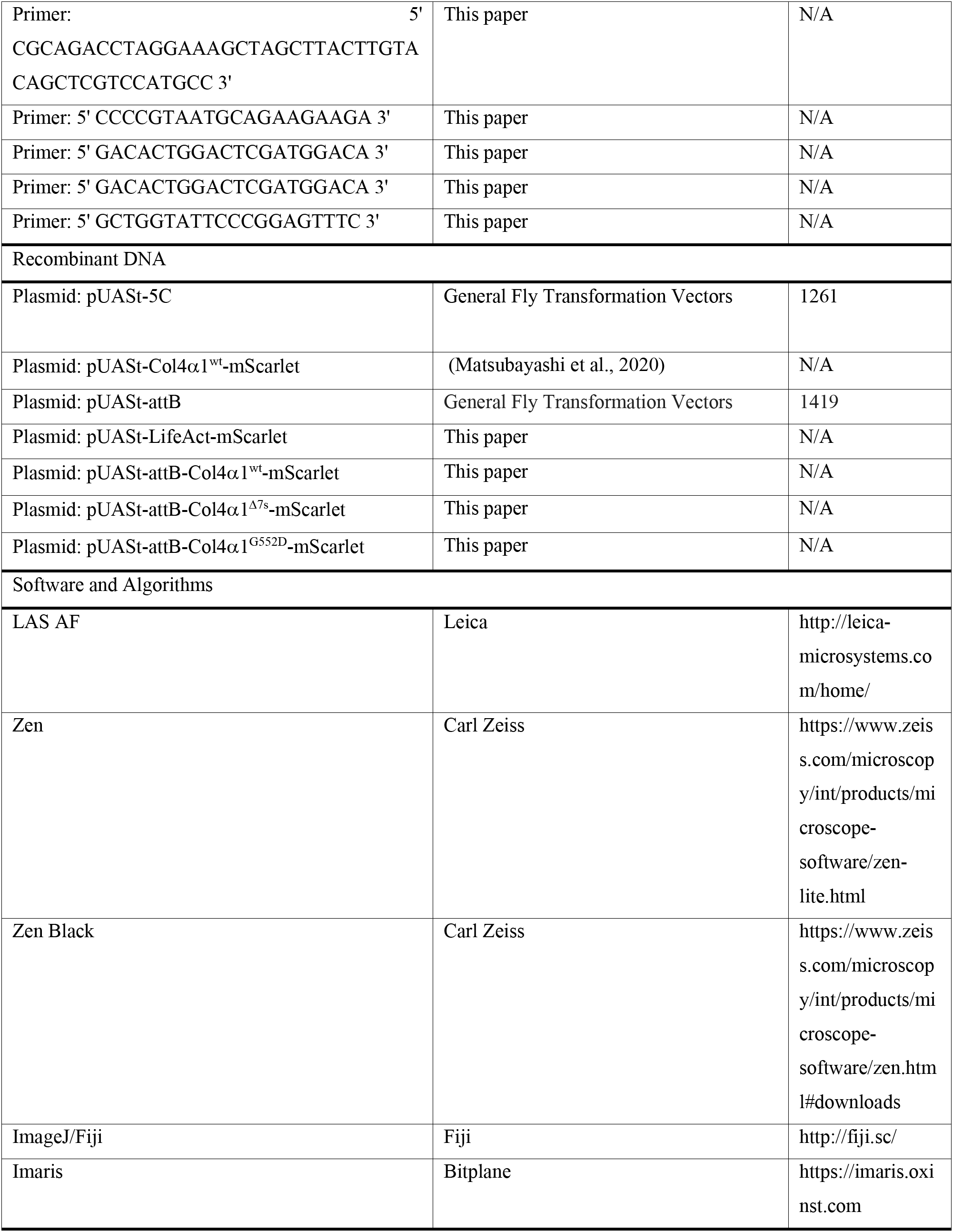
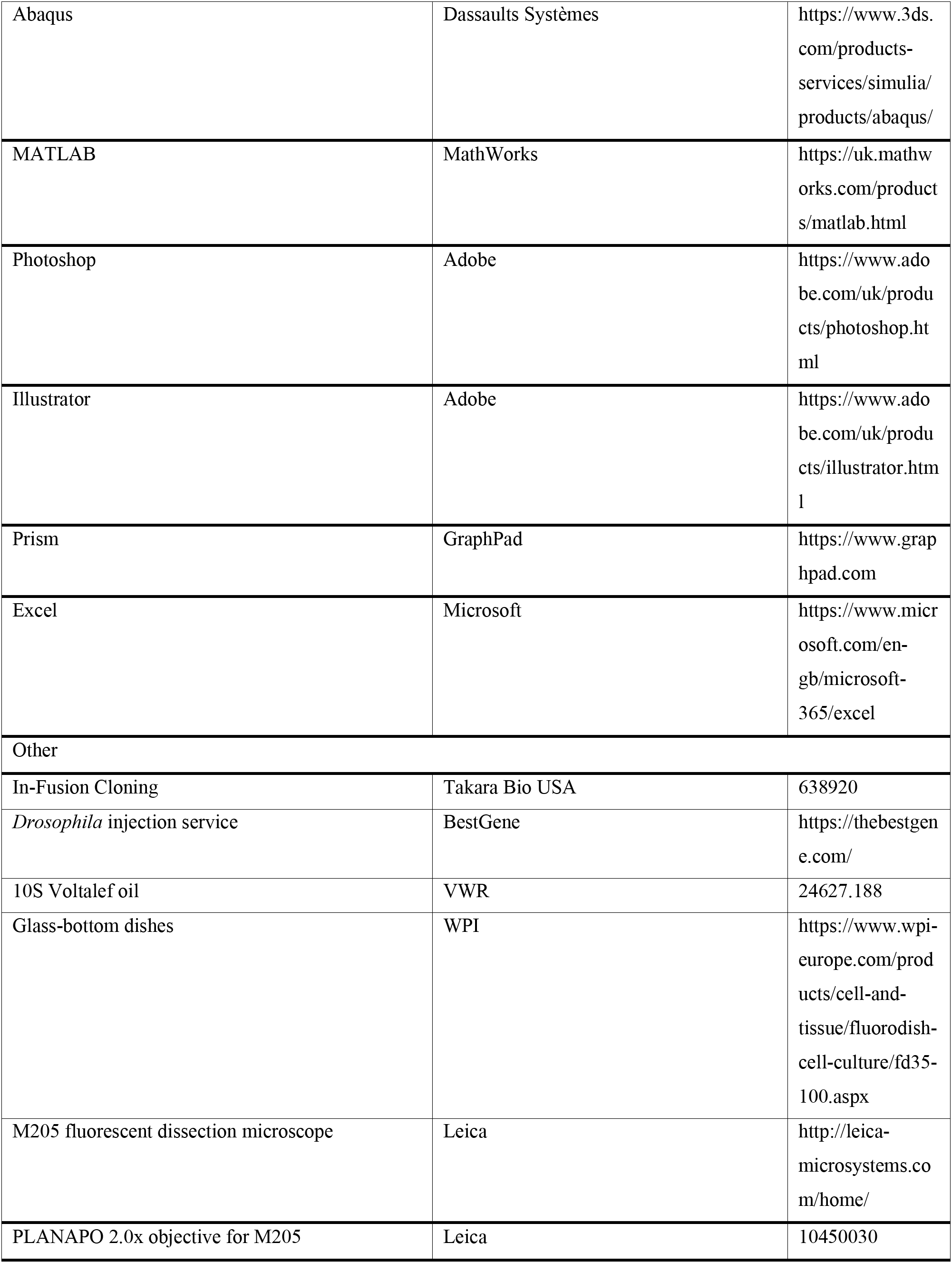
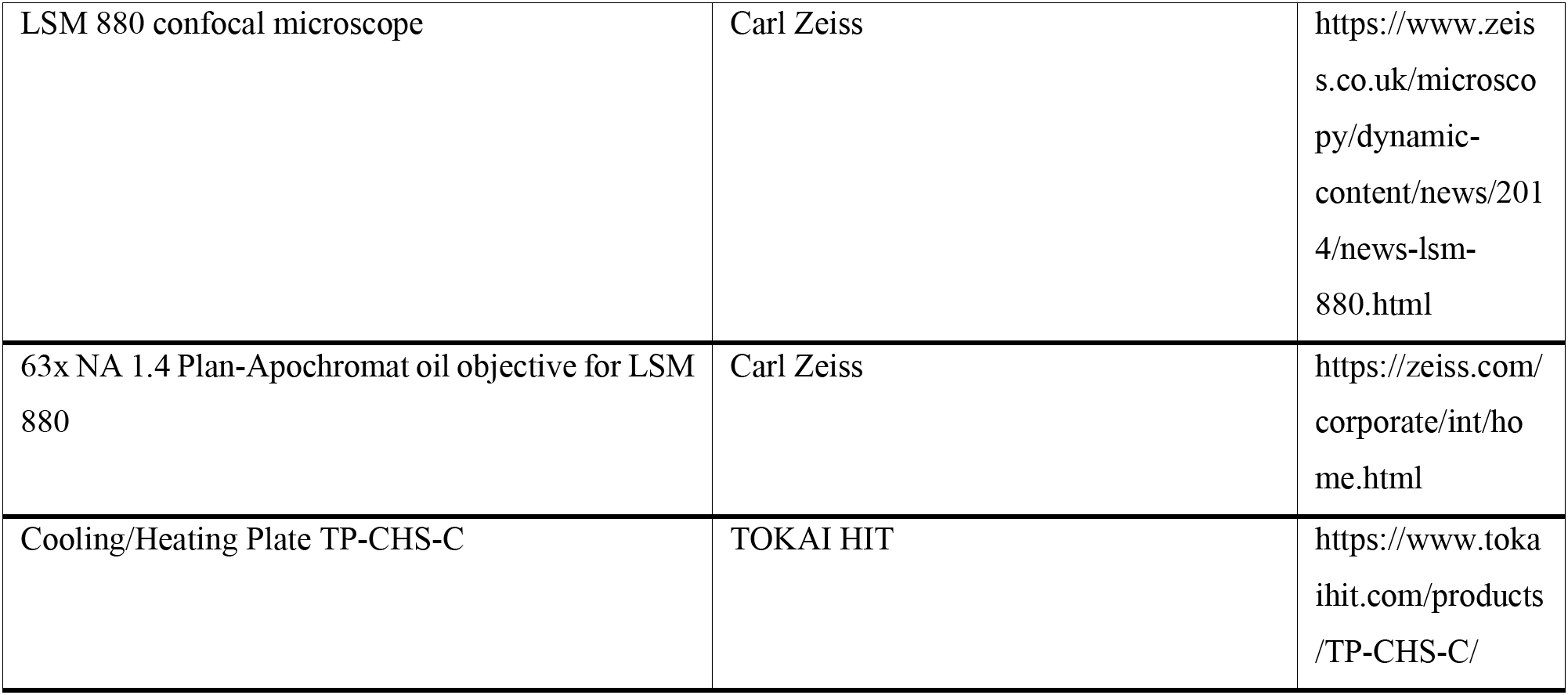

### Fly stocks and preparations

Repo-Gal4 (Sepp and Auld, 1999) and elav-Gal4 (Luo *et al*., 1994) were used to target transgene expression in the glial cells and neurons forming the VNC. Sn-Gal4 (Zanet *et al*., 2012) was used to express transgenes specifically in hemocytes. The parasegmental driver PS7-Gal4 (Bowman *et al*., 2014) was used to express transgenes in a narrow stripe along the VNC. The following UAS lines were used: UAS-LifeActGFP (Zanet *et al*., 2012), UAS-RedStinger, UAS-MMP2 (Page-McCaw *et al*., 2003), UAS-Rac DN (Luo *et al*., 1994), UAS-Myosin II DN (Franke *et al*., 2005), UAS-Col4α2-GFP (Van de Bor *et al*., 2015), UAS-LifeAct-mScarlet, UAS-Col4α1^wt^-mScarlet, UAS-Col4α1^Δ7S^-mScarlet, and UAS-Col4α1^G552D^-mScarlet (see following sections). Fax::GFP (Buszczak *et al*., 2007), repo-Cherry (gift from Clemens Cabernard, University of Basel), and elav-mYFP (Kohsaka *et al*., 2007) were used to label the VNC independently of Gal4. SrpHemo-H2A::3xmCherry (Gyoergy *et al*., 2018) was used to label the nuclei of hemocytes independently of Gal4. The homozygous viable Col4α2-GFP (Morin *et al*., 2001) protein trap was used to visualize Col4. The following mutant alleles and deficiencies were used: Df(2L)LanB1 (referred to as ΔLaminin, removing lanB1) (Urbano *et al*., 2009), Df(2L)BSC172 (referred to as ΔCol4, removing a chromosomal region including the two Drosophila Col IV genes vkg and Cg25C), Cg25C^k13420^ (Col4α1 mutant), trol^null^ (referred to as ΔPerlecan) (Voigt *et al*., 2002), srp^AS^ (lacking hemocytes) (Rehorn *et al*., 1996), Col4α1^G552D^ (referred as G552D and also known as *DTS-L2*, containing an aspartatic acid substitution in Col4α1 at G552) (Kelemen-Valkony *et al*., 2012), *Mhc*^1^ (gift from Frank Schnorrer and characterized for its use in imaging the *Drosophila* embryo in (Matsubayashi *et al*., 2020)). Unless used for temperature switch experiments, the flies were left to lay eggs on grape juice agar plates overnight at 25°C. Embryos were dechorionated in bleach. The appropriate genotype of the embryos was identified based on the presence of fluorescent probes and/or the absence of balancer chromosome expressing fluorescent markers. The genotypes of the embryos used in each experiment are listed in the next section.

### Genotypes of the embryos used in each experiment

Figure 1

A

*w; Mhc^1^; Repo-Gal4, UAS-LifeActGFP, UAS-RedStinger /+*

B-C

*w; Mhc^1^; Repo-Gal4, UAS-LifeActGFP /+*

D-E

*w; Col4α2-GFP; Repo-Gal4, UAS-LifeAct-mScarlet /+*

Figure 2

A

*“Control”: w;; Repo-Gal4, UAS-LifeActGFP/+*

*“ΔCol4”: w; Df(2L)BSC172; Repo-Gal4, UAS-LifeActGFP/+*

B

“Control”: *w;; Repo-Gal4, UAS-LifeActGFP /+*

“ΔPerlecan”: *w; trol^null^; Repo-Gal4, UAS-LifeActGFP /+*

“ΔCol4”: *w; Df(2L)BSC172; Repo-Gal4, UAS-LifeActGFP /+*

“ΔLaminin”: *w; Df(2L)LanB1; Repo-Gal4, UAS-LifeActGFP /+*

C-D

“Control”: *w;; repo-Gal4, UAS-LifeActGFP /+*

“ΔPerlecan”: *w; trol^null^; Repo-Gal4, UAS-LifeActGFP /+*

“ΔCol4”: *w; Df(2L)BSC172 / Cg25C^k13420^; Repo-Gal4, UAS-LifeActGFP /+*

“ΔLaminin”: *w; Df(2L)LanB1; Repo-Gal4, UAS-LifeActGFP /+*

Figure 3

B-E

*w; Mhc^1^, Col4α2-GFP; Repo-Gal4, UAS-LifeAct-mScarlet /+*

Figure 4

B

“Control”: *w;* PS7-Gal4 /+; faxGFP /+

“PS7>RacDN”: *w; PS7-Gal4 /+; faxGFP / UAS-RacN*^17^

“PS7>MMP2”: *w; PS7-Gal4 / UAS-MMP2; faxGFP /+*

C

“Control”: *w;; Sn-Gal4, UAS-Col4α2-GFP*

“G552D”: *w; Col4α1^G552D^; Sn-Gal4, UAS-Col4α2-GFP*

D

“G552D”: *w; Col4α1^G552D^; Sn-Gal4, UAS-Col4α2-GFP*

E

“Control”: *w;; repo-Gal4, UAS-LifeActGFP /+*

“G552D”: *w; Col4α1^G552D^; Repo-Gal4, UAS-LifeActGFP /+*

H

“Hemocyte>wt Col4α1”: *w; Mhc^1^, Col4α2-GFP; Sn-Gal4, UAS-Col4α1^wt^-mScarlet*

“Hemocyte>Δ7S Col4α1”: *w; Mhc^1^, Col4α2-GFP; Sn-Gal4, UAS-Col4α1^Δ7S^-mScarlet*

I

“Hemocyte>wt Col4α1”: *w; elav-mYFP; Sn-Gal4, UAS-Col4α1^wt^-mScarlet*

“Hemocyte>Δ7S Col4α1”: *w; elav-mYFP; Sn-Gal4, UAS-Col4α1^Δ7S^-mScarlet*

Supplementary Figure 1

A-B

*w; Mhc^1^; Repo-Gal4, UAS-LifeActGFP, UAS-RedStinger /+*

C

“Control”: *w;; Repo-Gal4, UAS-LifeActGFP /+*

“Glia>RacDN”: *w;; Repo-Gal4, UAS-LifeActGFP / UAS-RacN*^17^

D

“Control”: *w;; Repo-Gal4, UAS-LifeActGFP /+*

“Glia>Myosin II DN”: *w; Myosin II DN/+; Repo-Gal4, UAS-LifeActGFP /+*

E-F

*w;; Repo-Gal4, UAS-LifeActGFP, UAS-RedStinger / UAS-RacN*^17^

G

“Control”: *Elav-Gal4 /+;;Repo-mCD8::Cherry /+*

“Neuron>RacDN”: *Elav-Gal4 /+;;Repo-mCD8::Cherry / UAS-RacN*^17^

H

“Control”: *Elav-Gal4 /+;;Repo-mCD8::Cherry /+*

“Neuron>Myosin II DN”: *Elav-Gal4 /+; UAS-Myosin IIDN/+; Repo-mCD8::Cherry /+*

I-L

*Mhc^1^; w;; Repo-Gal4, UAS-LifeActGFP, UAS-RedStinger /+*

Supplementary Figure 2

A-B

*w; Mhc^1^, Col4α2-GFP*

C

“Control”: *w; Col4α2-GFP; Repo-Gal4, UAS-LifeAct-mScarlet /+*

“Glia>MMP2”: *w; Col4α2-GFP, UAS-MMP2 / Col4α2-GFP; Repo-Gal4, UAS-LifeAct-mScarlet /+*

D

“Control”: *w; Repo-Gal4, UAS-LifeAct-mScarlet /+*

“Glia>MMP2”: *w; UAS-MMP2 /+; Repo-Gal4, UAS-LifeAct-mScarlet /+*

“ΔCol4”: *w; Df(2L)BSC172; Repo-Gal4, UAS-LifeActGFP /+*

Supplementary Figure 3

A-D

“Control”: *w; elav-mYFP; Sn-Gal4, UAS-Col4α1^wt^-mScarlet*

“ΔHemocytes”: *w; SrpA^S^; elav-mYFP*

“Hemocytes>RacDN”: *w; elav-mYFP; Sn-Gal4, UAS-Col4α1^wt^-mScarlet / UAS-RacN*^17^

Supplementary Figure 4

A-C

*w; Mhc^1^, Col4α2-GFP; Repo-Gal4, UAS-LifeAct-mScarlet /+*

D-E

“Hemocytes>wt Col4α1”: *w; Mhc^1^, Col4α2-GFP; Sn-Gal4, UAS-Col4α1^wt^-mScarlet*

“Hemocytes>G552D Col4α1”: *w; Mhc^1^, Col4α2-GFP; Sn-Gal4, UAS-Col4α1^G552D^-mScarlet*

F

“Hemocytes>wt Col4α1”: *w; elav-mYFP; Sn-Gal4, UAS-Col4α1^wt^-mScarlet*

“Hemocytes>G552D Col4α1”: *w; elav-mYFP; Sn-Gal4, UAS-Col4α1^G552D^-mScarlet*

### Construction of UAS-LifeAct-mScarlet, UAS-Col4α1^wt^-mScarlet, UAS-Col4α1^Δ7s^-mScarlet and UAS-Col4α1^G552D^-mScarlet

<u>pUASt-LifeAct-mScarlet</u> was generated by inserting an 802 bp fragment (synthesized by eurofins Genomics) containing LifeAct followed by mScarlet-I sequences (Bindels et al., 2017), and an extra 15 bp at the 3’ and 5’ ends allowing their insertion into the linearized NheI-PacI pUASt-5C plasmid using In-Fusion cloning strategy (Takara Bio USA, Inc.).

The construct was sequenced (eurofins Genomics) using the following sequencing primers:

5’ GGGAATTGGCCGGCCAATTAATTAAATGGGTGTCGCAGATTTGATCAAG 3’
5’ CGCAGACCTAGGAAAGCTAGCTTACTTGTACAGCTCGTCCATGCC 3’

<u>pUASt-attB-Col4α1^wt^-mScarlet</u> was generated inserting a 15 kb fragment containing the Col4α1^wt^-mScarlet sequence from pUASt-Col4α1^wt^-mScarlet (Matsubayashi *et al*., 2020), into the linearized PacI-AvrII pUASt-attB plasmid using ligation T4 strategy (New England Biolabs, Inc.).

<u>pUASt-attB-Col4α1^Δ7s^-mScarlet</u> was generated by replacing a region of the 7s domain sequence of the pUASt-attB-Col4α1^wt^-mScarlet with a truncated version (Figure S4). A 950 bp fragment containing the 7s region was excised from the plasmid using XhoI and AsiSI sites and substituted by a 588 bp fragment (synthesized by Bio Basic USA, Inc.) containing the 7s truncated region, into the linearised XhoI-AsiSI pUASt-attB-Col4α1^wt^-mScarlet plasmid using ligation T4 strategy (New England Biolabs, Inc.).

The constructs were sequenced using the following sequencing primers:

5’ CCCCGTAATGCAGAAGAAGA 3’
5’ GACACTGGACTCGATGGACA 3’

<u>pUASt-attB-Col4α1^G552D^-mScarlet</u> was generated by replacing the sequence for the 552 Glycine of Col4α1^wt^ sequence (GGC) of the pUASt-attB-Col4α1^wt^-mScarlet for a new sequence which produces an Aspartic acid (GAC). A 2003 bp region containing the wild-type version of 552 Glycine was excised from the plasmid using AsiSI and EcoNI sites and substituted by a 2003 bp fragment (synthesised by Bio Basic USA, Inc.) containing the sequence for 552 Aspartic, into the linearized AsiSI-EcoNI pUASt-attB-Col4α1^wt^-mScarlet plasmid using ligation T4 strategy (New England Biolabs, Inc.).

The constructs were sequenced using the following sequencing primers:

5’ GACACTGGACTCGATGGACA 3’
5’ GCTGGTATTCCCGGAGTTTC 3’

The plasmids obtained were injected into flies (BDSC 9744) which contain an attP-9A insertion at 3R chromosome (89E11) by BestGene.

### Sample preparation and mounting for imaging

For all the confocal imaging, dechorionated embryos were mounted in a drop of Voltalef oil (VWR) between a glass coverslip covered with heptane glue and a gas-permeable Lumox culture dish (Sarstedt) as described previously (Davis et al., 2012). To image the embryos during stage 17 of embryogenesis on the confocal microscope, muscle myosin heavy chain (*Mhc*^1^) mutant embryos that do not twitch but have normal VNC condensation and expression of BM components were used as previously described (Matsubayashi *et al*., 2020). For the widefield imaging, the embryos were mounted in the same way but without heptane glue. Temperature controlled experiments were performed by mounting the gas-permeable Lumox culture dish on a TOKAI HIT TP-CHS-C heating/cooling plate.

For the AFM experiments, embryos were prepared as previously described (Matsubayashi et al., 2017), (Lee et al., 2009; Sanchez-Sanchez et al., 2017) After dechorianation with bleach, embryos were transferred to the heptane glue covered glass bottom of a 35 mm cell culture dish (FD35-100) which was then filled with 1x PBS. A cut was made along the dorsal midline of the embryos using an insect pin/needle, exposing the dorsal surface of the VNC.

### Widefield and confocal microscopy

Widefield images were acquired every 2 min with an M205 fluorescent dissection microscope (Leica) equipped with a PLANAPO 2.0x objective. All the confocal imaging was performed using an LSM880 confocal microscope (Carl Zeiss) equipped with a 63x NA 1.4 Plan-Apochromat oil objective unless stated otherwise. Tilescans (8 × 2 overlap 10%) of 40 μm Z-stacks with a zoom of 1.2 were used for PIV dynamics of the full VNC, with lateral view and temporal resolution of 10 min. For live imaging of Col4 and glia in the tail of the VNC, 20 μm Z-stacks of Col4α2-GFP and repo>LifeAct-mScarlet were acquired with a 63X objective with 2x zoom, and a temporal resolution of 5 min/frame using Optimal Airyscan mode. For simultaneous imaging of Col4 in head vs. tail of the VNC, multipoint acquisition consisting of 20 μm Z-stacks of Col4α2-GFP in head and tail regions of the same embryo were acquired with a 63X objective with 4x zoom, and a temporal resolution of 30 sec/frame using Super Resolution Airyscan mode. To visualize Col4 on the surface of the full VNC with a ventral view, tilescans (10 × 2 overlap 0%) of 20 μm Z-stacks were acquired on Super Resolution Airyscan mode with a zoom of 4. For volume measurements of the full VNC, two tiled images (overlap 10%) of 100 μm Z-stacks with a 20x air objective (Plan-Apochromat air objective, NA 0.8) were acquired with a temporal resolution of 10 min. Images were stitched using the Zen Black software and exported to the Imaris software (Bitplane) for further analysis.

### VNC condensation and Col4 levels quantification by fluorescent dissection microscope

For the quantification of VNC condensation, the last position of the tail of the VNC was manually tracked on every frame (2 min time resolution) by using Fiji (Schindelin et al., 2012). The x and y coordinates of each point were stored and compared to the initial position of the VNC tail to quantify displacement through time.

For the quantification of relative Col4α2-GFP-trap levels through time (2 min time resolution), the average raw fluorescence intensity of each embryo in every frame was measured with Fiji, the acquired data were smoothed by calculating a 10-frame moving average.

### Volume and surface measurements

To visualize the VNC and measure its volume, the glial cells of the VNC were labelled with UAS-LifeAct-mScarlet under the expression of the glia driver repo-Gal4. To determine the 3 timepoints relevant to the condensation process we proceeded as follows: the beginning of the 1^st^ phase of condensation was determined as the first frame in which displacement of the VNC tail was observed; the end of the 1^st^ phase was determined as 3 hours from the beginning of the 1^st^ phase; and the end of the 2^nd^ phase as 9 hours from the end of the 1^st^ phase. After the frames were selected, each peripheral nerve sprouting from the VNC was manually deleted, by using the “Surface” function in Imaris and by turning the voxels from the magenta channel into zero in the selected areas. Subsequently, the volume of the VNC was obtained by generating a surface on the magenta channel with a voxel of size 2. The length, width, and height of the VNC were quantified with the Imaris function “Measurement Points”. For each embryo, the average height and width were measured at the central point of each neuromere of the VNC. We used the measured VNC dimensions to calculate the change in surface area, using an idealized shape resembling an elliptical cylinder.

### Quantification of fluorescence intensity in the surface of the VNC

To quantify the Col4 gradient on the VNC surface from confocal images, we first removed the signal from the intra-cellular Col4α2-GFP inside hemocytes by discarding any signal higher than 70% of the total fluorescence intensity. We then calculated the average signal across the VNC from head to tail smoothed with a walking average over 200 μm.

### Particle image velocimetry (PIV) analysis

PIV was performed with a custom MATLAB suite (https://github.com/stemarcotti/PIV) as in (Davis et al., 2015; Yolland et al., 2019). This methodology was applied on time series confocal imaging of the VNC and was used to track the motion on the VNC surface over time. Briefly, the algorithm cross-correlates a small region of interest in a frame of reference (source area) to a larger portion of the subsequent time frame (search area), to find the best match. This operation is performed iteratively on all portions of the image at each time frame to allow for feature tracking in the entire field of view. The parameters were optimized as follows: source size 9 μm, search size 16 μm, grid size 5 μm, correlation threshold 0.5, when the entire VNC was imaged; source size 2 μm, search size 4 μm, grid size 1 μm, correlation threshold 0.3, when only a portion of the VNC was imaged with high spatiotemporal resolution.

The obtained displacement vector field was then interpolated both spatially and temporally. The interpolation kernel parameters were set as follows: spatial kernel size 50 μm (sigma 10 μm), temporal kernel size 90 min (sigma 40 min), when the entire VNC was imaged; spatial kernel size 10 μm (sigma 2 μm), temporal kernel size 5 frames (sigma 2 frames), when only a portion of the VNC was imaged with high spatiotemporal resolution. All colourmaps presented were generated to visualize the magnitude of feature velocity, and vectors from the interpolated field were overlaid to highlight flow direction. An average of the velocity magnitude or x component across the length of the VNC was calculated for each frame to plot kymographs.

### PIV local alignment correlation

To evaluate the coherence of motion for collagen and glia, 256 random locations were selected on the non-interpolated PIV displacement vector field for each time frame. The displacement vector at each of these locations was considered as a reference, and its orientation was compared to all the neighboring vectors over an 8 μm radius. This was done by calculating the norm of the cosine of the angle theta between the reference vector and each of the neighboring vectors, and by subsequently obtaining an average across all these comparisons for each location. The computed cosine values were smoothed with a walking average over 5 frames (25 min). Values close to 1 indicate similar orientation between the reference and its neighbors, representing high motion coherence.

A similar approach was taken to measure the coherence of motion for collagen in the tail-to-head direction for two distinct locations at the head and tail of the VNC. For each frame, the direction of 11×11 vectors over a 3 μm-spaced grid of the interpolated PIV field was compared to the reference direction, i.e., tail-to-head direction. The cosine of the angle theta between each vector and the reference vector was calculated as a measure of coherence of motion in the tail-to-head direction, and an average calculated for each frame. Values close to 1 indicate motion in the tail-to-head direction.

### Cell tracking

Confocal time-lapse movies of the glial cells forming the VNC containing labelled nuclei were tracked with the Imaris function “Spots”. The tracking was divided in two areas, anterior (head) and posterior (tail), which were defined as half of the VNC at the end of the initial 3 hours of VNC condensation. Two 3-hour periods were chosen in the first and second phase, respectively (0-3 hours and 7-10 hours from the start of VNC condensation).

The obtained tracks were analyzed with custom software in MATLAB to obtain information on their directionality. The start and end (x,y) coordinates of each track within these time periods which was longer than 10 frames (100 min) were extracted. A vector was drawn between each start and end location and its angle computed to display the polar histograms (angles corresponding to zero were set on the horizontal axis from head to tail). 81 and 130 tracks were analyzed for the first phase for head and tail, respectively; 219 and 206 tracks were analyzed for the second phase for head and tail, respectively. The mean average velocity of all the tracks with a minimum length of 10 frames (100 min) in both areas was obtained from the Imaris function “Spots”.

### Atomic force microscopy (AFM)

Spherical tipped cantilevers were produced by gluing a 10 μm silica bead at the extremity of qp-CONT probes (Nanosensors). The cantilever spring constant was calibrated in liquid by thermal noise method (0.2 N/m). Measurements were performed using a custom-built system(Hirvonen et al., 2020), based on a standard inverted microscope (Axio Observer.Z1, Zeiss) and equipped with an atomic force microscope (NanoWizard 3, JPK Instruments).

Force spectroscopy measurements using a 3 nN set point were performed on the VNC towards the head side of the tissue. Analysis of the force spectroscopy curves to obtain the effective modulus was carried out in MATLAB by using custom-made algorithms as described in (Marcotti et al., 2019), considering a target indentation depth for the Hertz model fitting of 230 nm.

### Statistics and reproducibility

Statistical tests employed are listed in the caption of relevant figure panels. Significance was indicated as follows: ‘****’ for p-values lower than 0.0001, ‘***’ lower than 0.001, ‘**’ lower than 0.01, ‘*’ lower than 0.05, ‘ns’ otherwise.

## Supplementary material

## Appendix

### 1 Introduction

In this document, we illustrate in further detail the Finite Element (FE) model simulations of ventral nerve cord (VNC) condensation presented in the main text. Information on how the model was constructed, on how simulations were run and on parameters choice are given.

VNC morphogenesis involves two distinct phases, a first rapid isovolumetric phase lasting about 3 hours which shows an anisotropic 25% reduction in the length of the tissue from tail to head, and a second slower phase which shows a more isotropic condensation and a reduction in volume. As most of the morphological changes happen in the first phase of condensation, the modelling approach presented here focuses on this part of the process, with the aim of explaining the loading conditions necessary to initiate the anisotropic motion.

All FE simulations were performed in Abaqus (Simulia, Dassault Systèmes) by using static modelling. An idealised geometry was employed with dimensions acquired through experimental observations (Fig. 1C in the main text) and different loading conditions were simulated.

### 2 FE model

An idealised shape representing the approximate VNC morphology was designed as an elliptical cylinder (Fig. 1). The head side of the VNC is physically connected to the embryonic brain, while the tail of the tissue is free to move and has a semi-spherical end. Relevant dimensions were chosen according to experimental observations (Fig. 1C in the main text), with the elliptical cylinder measuring 400 *μm* in length, and displaying a cross section of major and minor axes measuring 83 *μm* and 38 *μm*, respectively.

**Figure 1:**
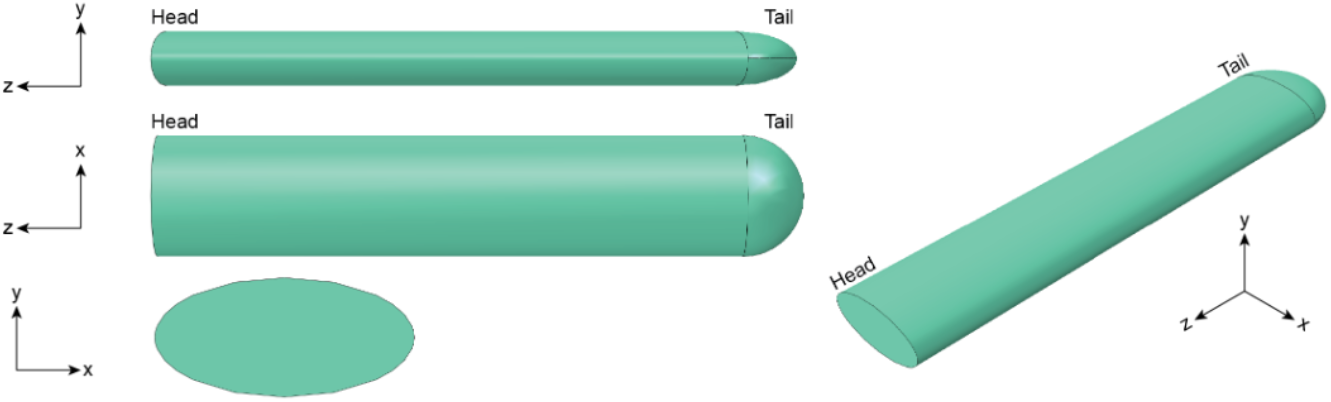
Geometry. Orthogonal views of the FE model of the VNC. The model is constituted by an elliptical cylindrical shape which dimensions were selected to match the experimental observations. The head side is connected to the tissue forming the brain, while the tail end is free to move.

The model was constituted by two materials, a tissue core which represents glia and neurons, and a surrounding thin shell representing the basement membrane. This surrounding layer was modelled as a 0.1 *μm*-thick skin shell. Both materials were defined as linear elastic with Poisson ratio of 0.48. Default Young’s moduli were set to 70 *Pa* for the tissue and 30 *Pa* for the basement membrane, if not otherwise specified in certain simulation cases. These values were chosen to match the experimental observations performed by atomic force microscopy (Fig. 2A in the main text). A shrinking coefficient was assigned to the shell layer when simulating surface tension.

To perform FE analysis, the geometry has to be meshed into a finite number of elements of known dimension. The geometry was therefore meshed with a total of 79818 nodes and 61689 elements for analysis, corresponding to elements of about 5 *μm* in size (Fig. 2). Two different types of elements were used: 7436 linear triangular elements for the shell (type S3) and 54253 quadratic tetrahedral elements for the tissue (type C3D10). Convergence tests were run for a range of different element dimensions (from 2.5 to 20 *μm*), each analysis showed little variation in the simulated reduction in length when loading the model with uniform pressure.

**Figure 2:**
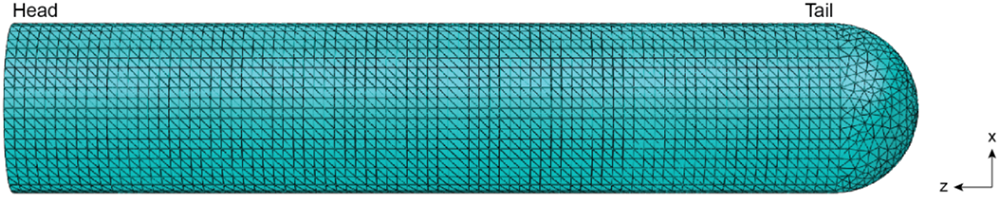
Mesh. Top-down view of the FE model showing the mesh used, with elements of about 5 *μm* in size. Linear triangular elements or quadratic tetrahedral elements were employed for the shell and tissue, respectively.

The model was constrained at the head side, where the VNC is attached to the brain tissue in the *Drosophila* embryo. As the brain tissue is of considerable dimensions compared to the VNC, it was hypothesised that this link could be simulated with an encastre, constraining all translational and rotational degrees of motion (Fig. 3).

**Figure 3:**
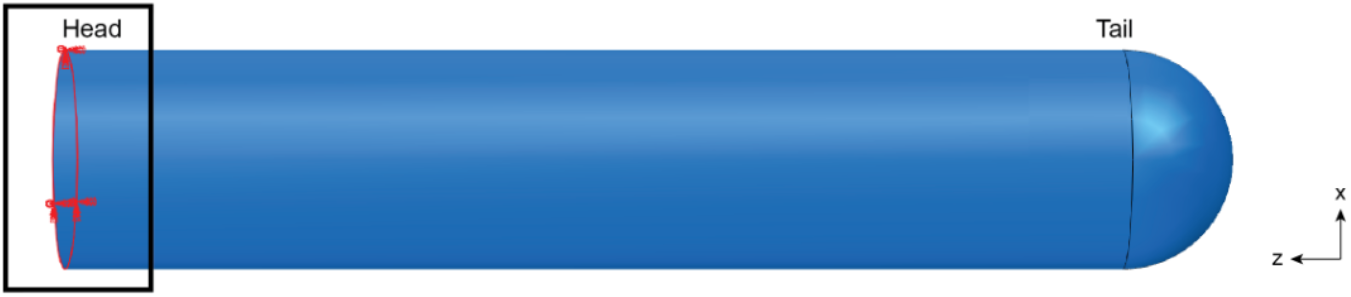
Boundary conditions. The model was constrained by imposing an encastre at the head side (black square) simulating its attachment to the brain tissue in the *Drosophila* embryo.

### 3 Loading scenarios

Different loading scenarios were simulated with the goal of reproducing the experimentally observed change in shape of the VNC during the first phase of morphogenesis. This phase of the process is characterised by an anisotropic reduction in length from the tail to the head of the tissue. The first phase does not display a change in the volume of the VNC, and the cross-section of the tissue expands along both axes to compensate for the reduction in length (Fig. 1C in the main text).

The different simulations performed and their rationales are outlined in the following sections. The simulated magnitude of the displacement along the length of the VNC was used as output, as a proxy for the experimentally observed motion of the tissue.

#### 3.1 Uniform pressure

We first loaded the model with a uniform pressure, normal to each element. This would correspond to a cell-driven active compression applied on the tissue, as the cells would locally contract resulting in a perpendicular force on the VNC. This loading scenario caused a reduction in length from tail to head as experimentally observed; however, it was accompanied by a reduction in the size of both axes in the cross-section, suggesting a change in volume which diverged from experimental data (Fig. 4). Increasing the magnitude of the applied pressure resulted in a more marked shrinking in all axes (Fig. 5). To obtain a reduction in length of 25%, a uniform pressure of 0.38 *kPa* was applied.

**Figure 4:**
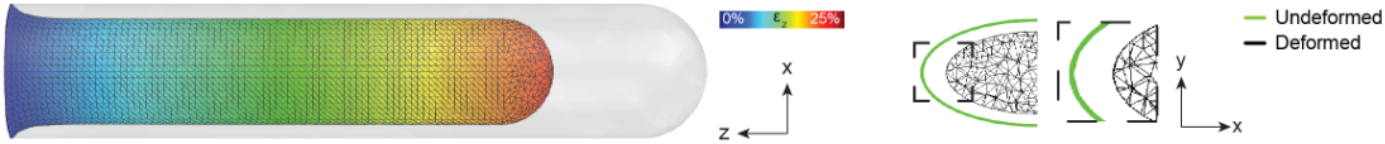
Uniform pressure. Simulating loading with a perpendicular uniform pressure on the VNC resulted in its anisotropic reduction in length from tail to head (left panel). However, reduction in both the height and width of the cross-section was also observed (right panel), diverging from experimental evidence. Colours represent the magnitude of displacement along the length of the tissue (z-axis) with values ranging from 0% to 25% of strain.

**Figure 5:**
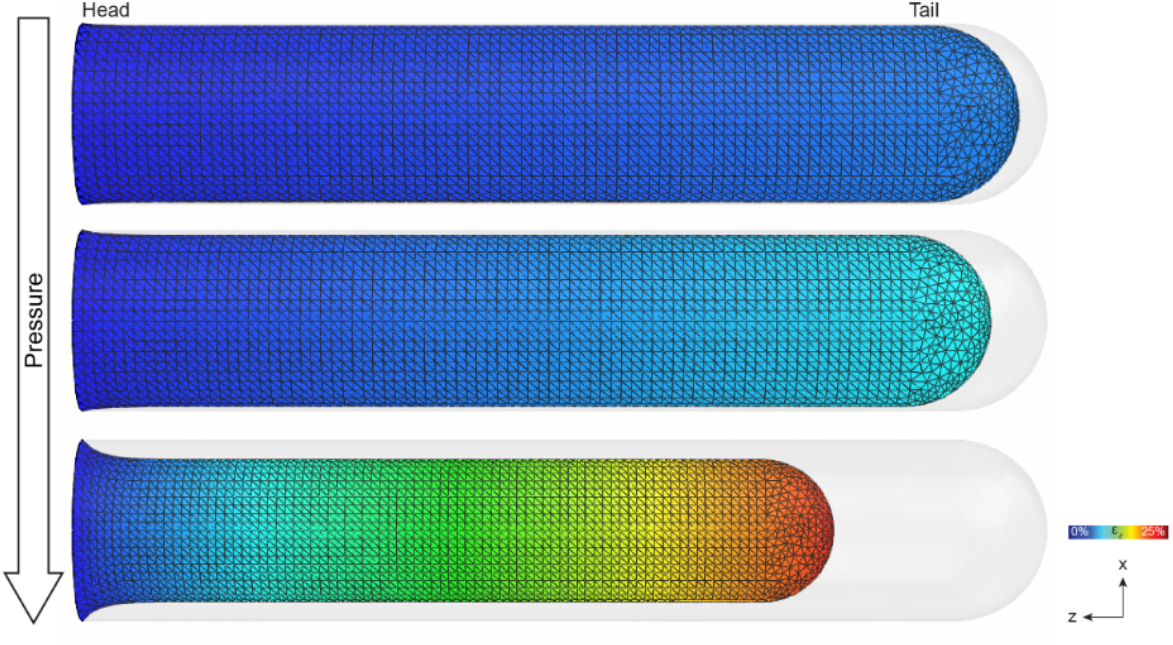
Uniform pressure - pressure magnitude. Increasing the magnitude of the simulated normal pressure exacerbated the amount of length reduction (top panel: pressure magnitude 0.05 *kPa*, middle: 0.1 *kPa*, bottom: 0.38 *kPa*). Colours represent the magnitude of displacement along the length of the tissue (z-axis) with values ranging from 0% to 25% of strain.

#### 3.2 Isotropic surface tension

We then simulated a loading scenario involving surface tension, as we hypothesised a role for the basement membrane layer surrounding the tissue in driving its condensation. To this aim, we assigned a temperature-controlled expansion coefficient to the shell material: by simulating a drop in temperature, we could cause the material of the shell to shrink. Similarly to loading the model with uniform pressure, this simulation resulted in an anisotropic reduction in length of the VNC. When looking at the cross-section of the tissue, we observed the expected increase in size along the y-axis, but also a decrease along the x-axis, with a tendency towards a more circular shape, which did not match with the experimental observations (Fig. 6).

**Figure 6:**
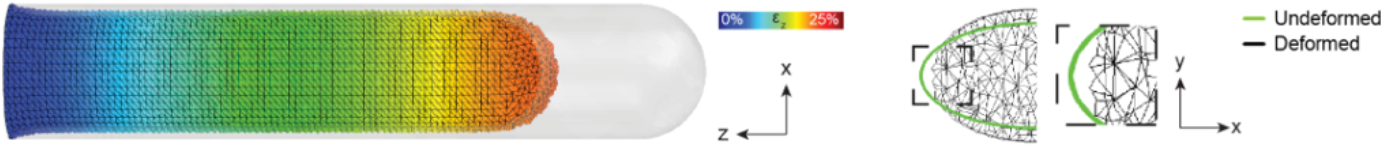
Isotropic surface tension. Simulating surface tension by allowing the shell material to shrink resulted in an anisotropic reduction in length towards the head (left panel). The change in shape in the cross-section showed increased dimensions in height and decreased dimensions in width (right panel). Colours represent the magnitude of displacement along the length of the tissue (z-axis) with values ranging from 0% to 25% of strain.

We could modulate the ability of the shell to shrink by keeping the temperature drop constant (i.e., 100 to 0 degrees) and subsequently changing the expansion coefficient (Fig. 7). This resulted in a progressively larger reduction in the VNC length. To simulate a 25% reduction in length, an expansion coefficient of 0.65 was required.

**Figure 7:**
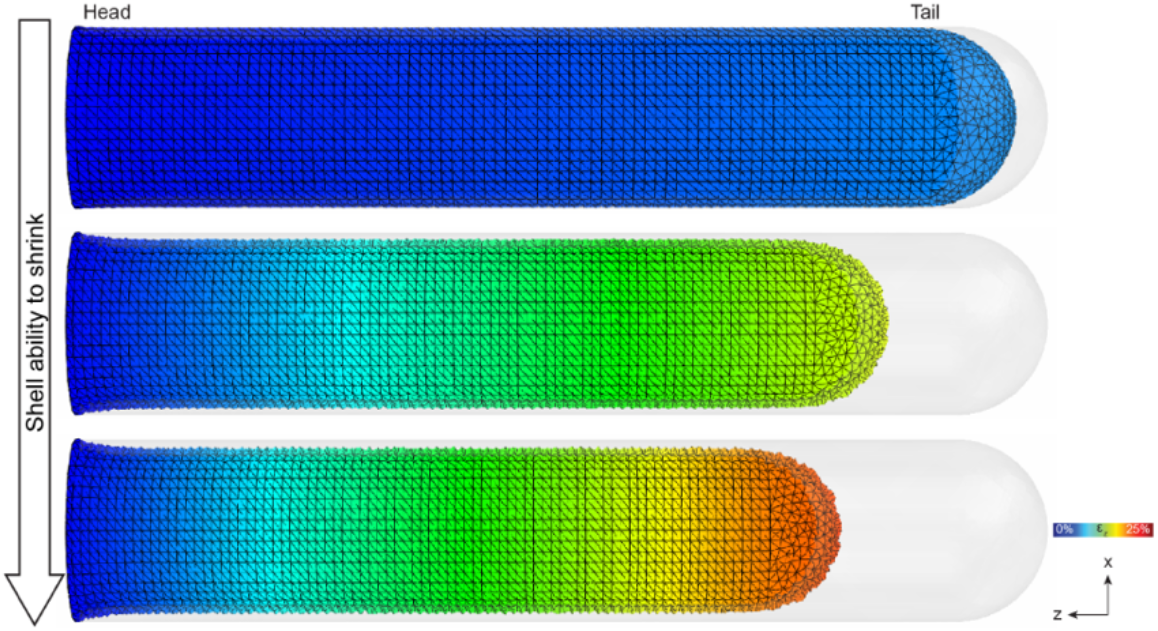
Isotropic surface tension - ability to shrink. Simulating surface tension by allowing the shell material to shrink resulted in an anisotropic reduction in length towards the head, proportional to the assigned expansion coefficient (top panel: expansion coefficient 0.1, middle: 0.5, bottom: 0.65). Colours represent the magnitude of displacement along the length of the tissue (z-axis) with values ranging from 0% to 25% of strain.

**Figure 8:**
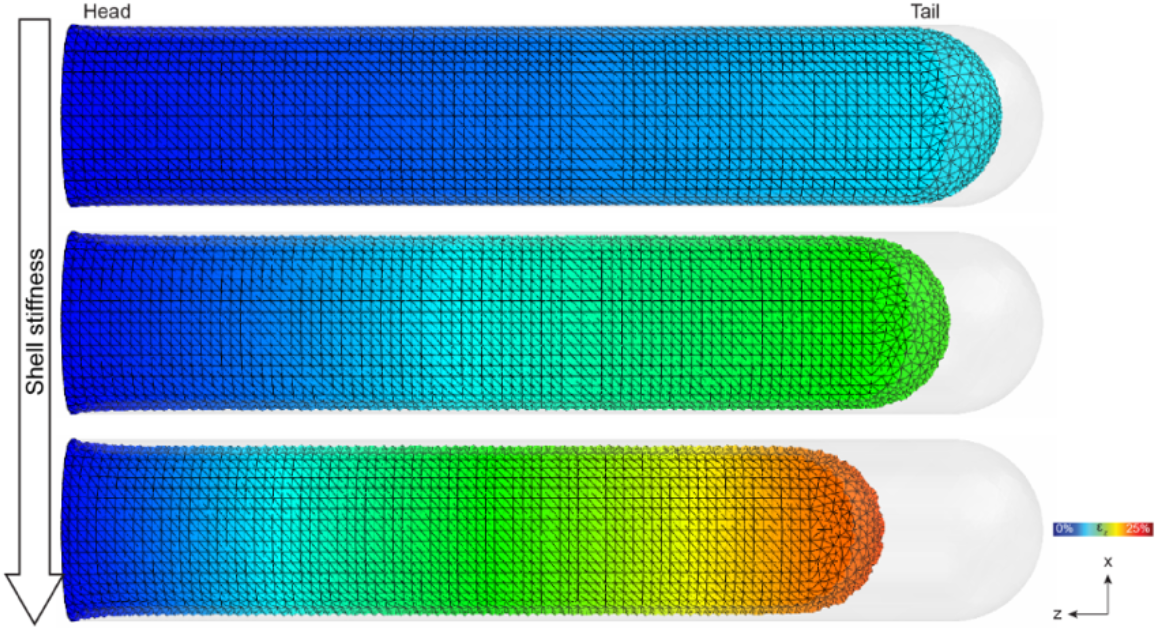
Isotropic surface tension - stiffness. Simulating surface tension by allowing the shell material to shrink resulted in an anisotropic reduction in length towards the head, proportional to the assigned shell stiffness (top panel: shell/tissue Young’s moduli 10/90 *kPa*, middle: 20/80 *kPa*, bottom: 30/70 *kPa*). Colours represent the magnitude of displacement along the length of the tissue (z-axis) with values ranging from 0% to 25% of strain.

**Figure 9:**
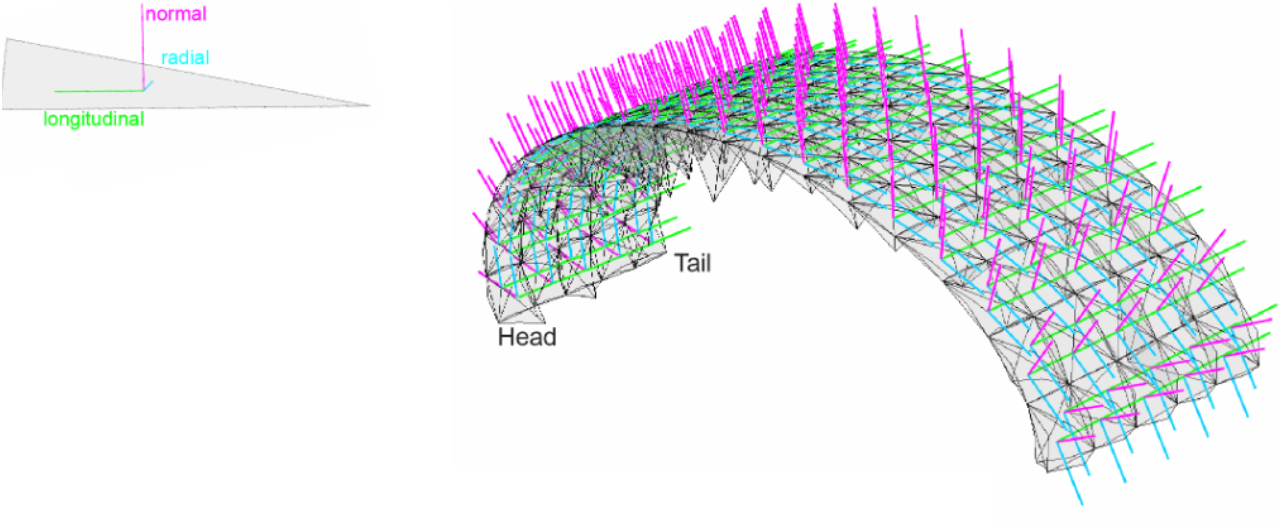
Material orientation. Simulating an anisotropic surface tension required assigning an expansion coefficient for each material orientation. Each element is assigned its own reference, with a normal (magenta), radial (cyan), and longitudinal (green) orientation.

The magnitude of the surface tension could also be adjusted by changing the material properties of the shell. In fact, by increasing the stiffness of the shell, it was possible to increase the reduction in length of the tissue, while keeping constant both the temperature drop (100 to 0 degrees) and expansion coefficient (0.65).

#### 3.3 Longitudinal surface tension

Finally, we simulated a longitudinal surface tension along the length of the tissue (z-axis), as we hypothesised that the observed gradient of collagen from head to tail may lead to a preferential directionality of surface tension (Supplementary Fig. 2AB). This was achieved by defining an anisotropic shell shrinking behaviour, which was described by an expansion coefficient for each of the three material orientations (9). We set to zero the expansion coefficients for the normal and radial orientations, and only kept active the longitudinal component for each element (0.65), which prevented the tissue from shrinking in the normal and radial axes. This showed an anisotropic reduction in VNC length from tail to head, which correlated with an expansion in both axes of the cross-section (10). This result was analogous to what was observed experimentally, suggesting that a longitudinal surface tension dictated by the collagen layer surrounding the tissue might be the driving force of the first phase of the condensation process.

#### 3.4 Comparison

While all three simulations (uniform pressure, isotropic surface tension, and longitudinal surface tension) showed an anisotropic reduction in the length of the tissue from tail to head (Fig. 11), the results when looking at the tissue cross-section varied, with only the longitudinal surface tension scenario predicting the expansion in both height and width of the tissue which was observed experimentally in the *Drosophila* embryo (Fig. 10).

**Figure 10:**
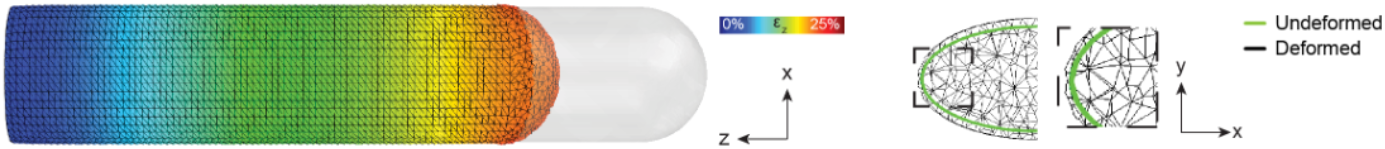
Longitudinal surface tension. Simulating a longitudinal surface tension by allowing the shell material to shrink only along the length of the tissue resulted in an anisotropic reduction in length towards the head (left panel) and an increase in both height and width of the tissue (right panel). This result mimicked the experimental observations in the *Drosophila* embryo. Colours represent the magnitude of displacement along the length of the tissue (z-axis) with values ranging from 0% to 25% of strain.

**Figure 11:**
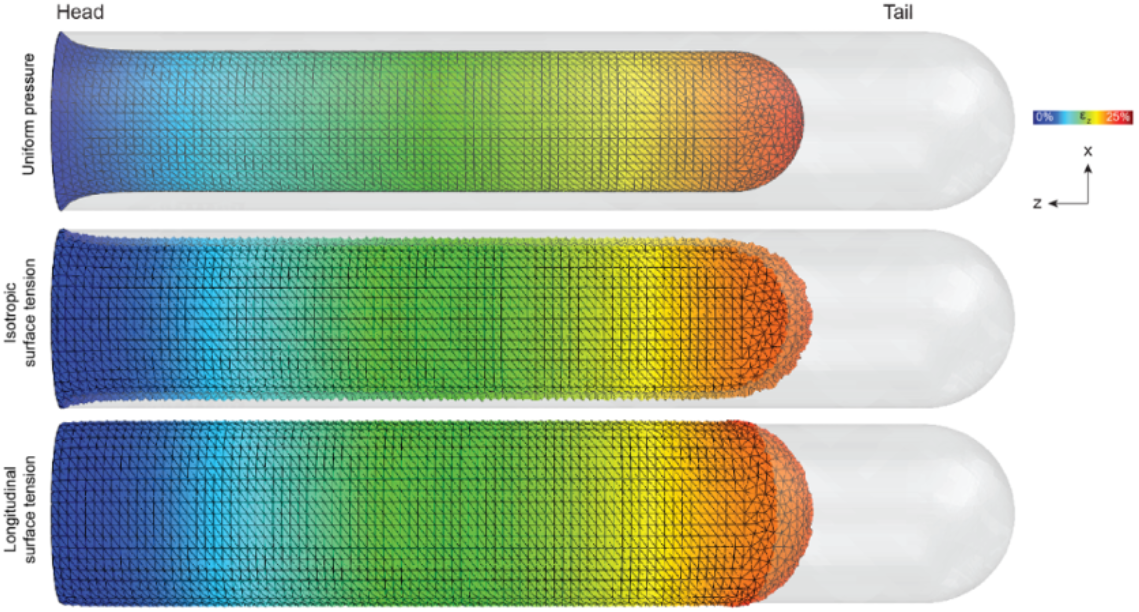
Displacement along the length of the tissue. Loading the model by uniform pressure, isotropic, and longitudinal surface tension resulted in an anisotropic reduction in the VNC length from tail to head. Colours represent the magnitude of displacement along the length of the tissue (z-axis) with values ranging from 0% to 25% of strain.

This difference is further highlighted by the vectorial representation of the displacement (Fig. 12): it can be noted how, in the case of the uniform pressure loading, the vectors point towards a central node at the head of the tissue suggesting a deformation along all axes. For the isotropic surface tension, this behaviour is less pronounced, but can still be observed towards the head showing additional deformation along the cross-section of the tissue. Finally, in the longitudinal surface tension case, the vectors have a tendency to point outwards from the tissue centre which explains the expansion in both x- and y-axis. The FE simulations, therefore, suggested that VNC condensation can be explained by a preferentially directed surface tension.

**Figure 12:**
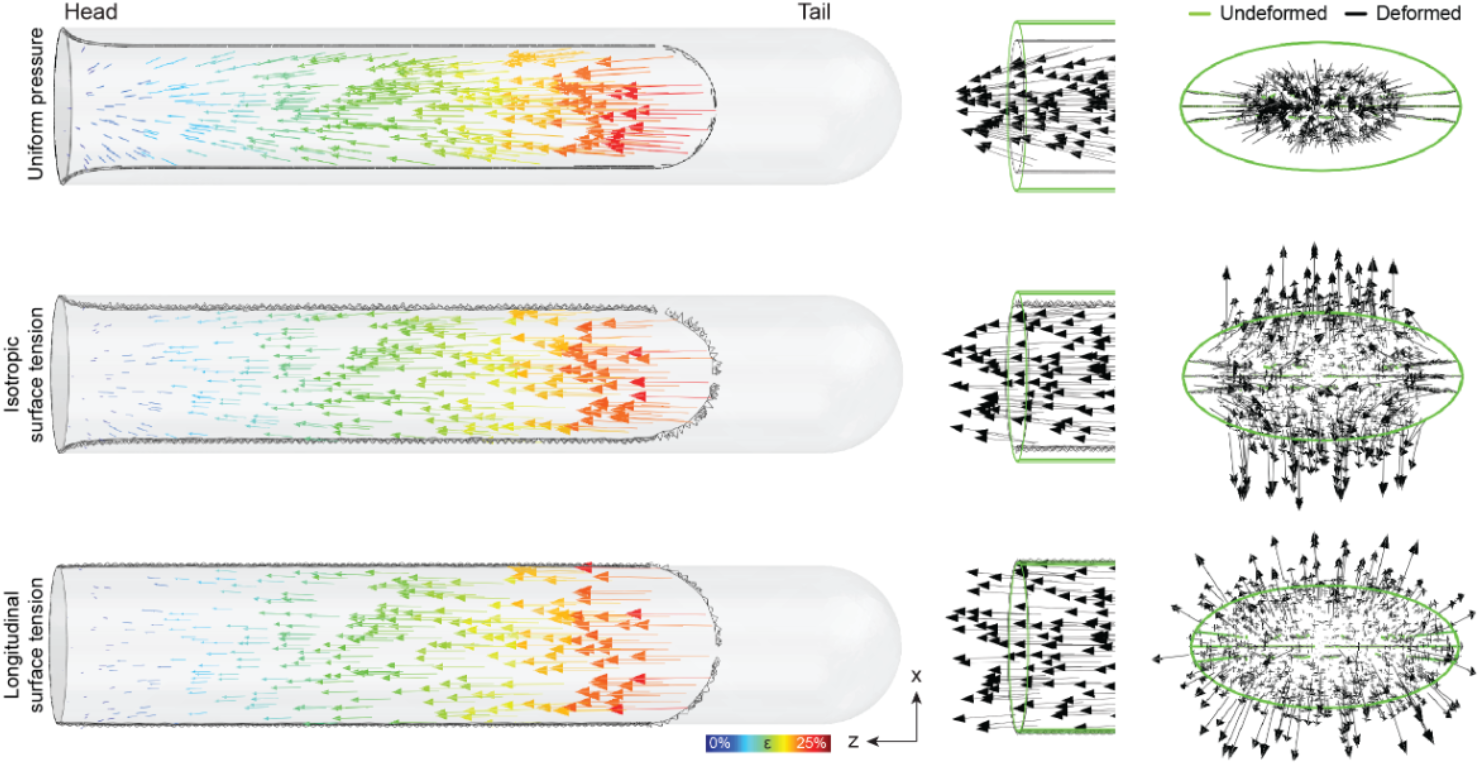
Displacement vectors. Vectorial representation of the displacement of the VNC when comparing loading the model by uniform pressure, isotropic, and longitudinal surface tension (left panels). The representation with unit vectors on a vertical cut along the VNC length highlights the directionality of the displacement (right panels). Colours represent the magnitude of overall displacement with values ranging from 0% to 25% of strain.

### 4 Disrupting the longitudinal surface tension

We hypothesised that an increase in longitudinal surface tension would drive the tissue reduction in length by global transmission of ECM-generated stresses through the polymer network. We therefore tested the effects of a local disruption in the surface tension on VNC condensation.

To achieve this, we defined the initial temperature field controlling the shell shrinking as a function of the tissue length (z-axis), before turning the temperature to 0 degrees to enable the shell shrinking. If no disruption in surface tension was to be simulated, the temperature value was kept constant along z (top panel Fig. 13, and Fig. 10). Otherwise, we analytically defined the shape of the initial temperature field along z with a Gaussian-shaped drop in temperature around the centre of the VNC (middle and bottom panels Fig. 13). The drop could be modulated to simulate a partial or null surface tension in a stripe of the tissue, by setting the target temperature to 50% (middle panel Fig. 13) or 0% (bottom panel Fig. 13) of the temperature away from the stripe.

**Figure 13:**
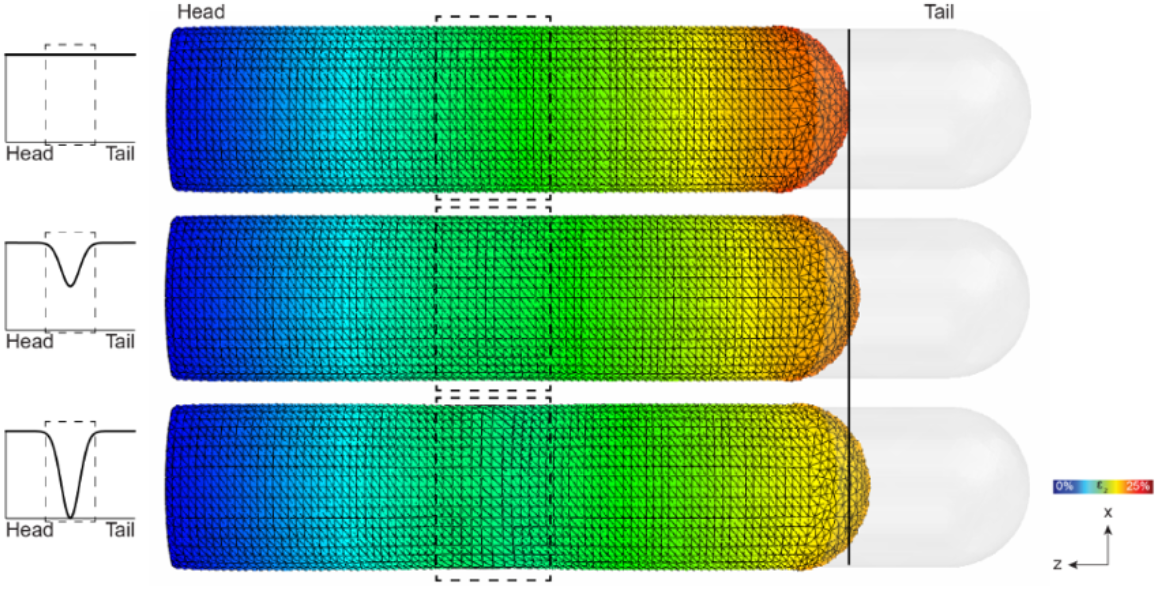
Disrupting surface tension. The reduction in VNC length was affected by locally disrupting the surface tension (black vertical line). This was achieved by locally modifying the initial temperature field of the simulation with a drop around the middle of the tissue (black squares) of either 50% (middle panel) or 100% (bottom panel), corresponding to halving or deleting the surface tension in a stripe of the VNC, respectively. Colours represent the magnitude of displacement along the length of the tissue (z-axis) with values ranging from 0% to 25% of strain.

Simulations revealed that a local disruption of the surface tension had an effect on the reduction in length of the tissue, proportional to the severity of the disruption (Fig. 13). Similar results were obtained when experimentally causing a local disruption of the collagen network, by expressing surface-bound matrix metalloproteases in a stripe along the VNC (Fig. 4B in the main text). These results taken together suggest that surface tension-driven change in tissue shape should involve long-range and coordinated morphological remodelling.

## Supplementary Figures

**Figure S1.**
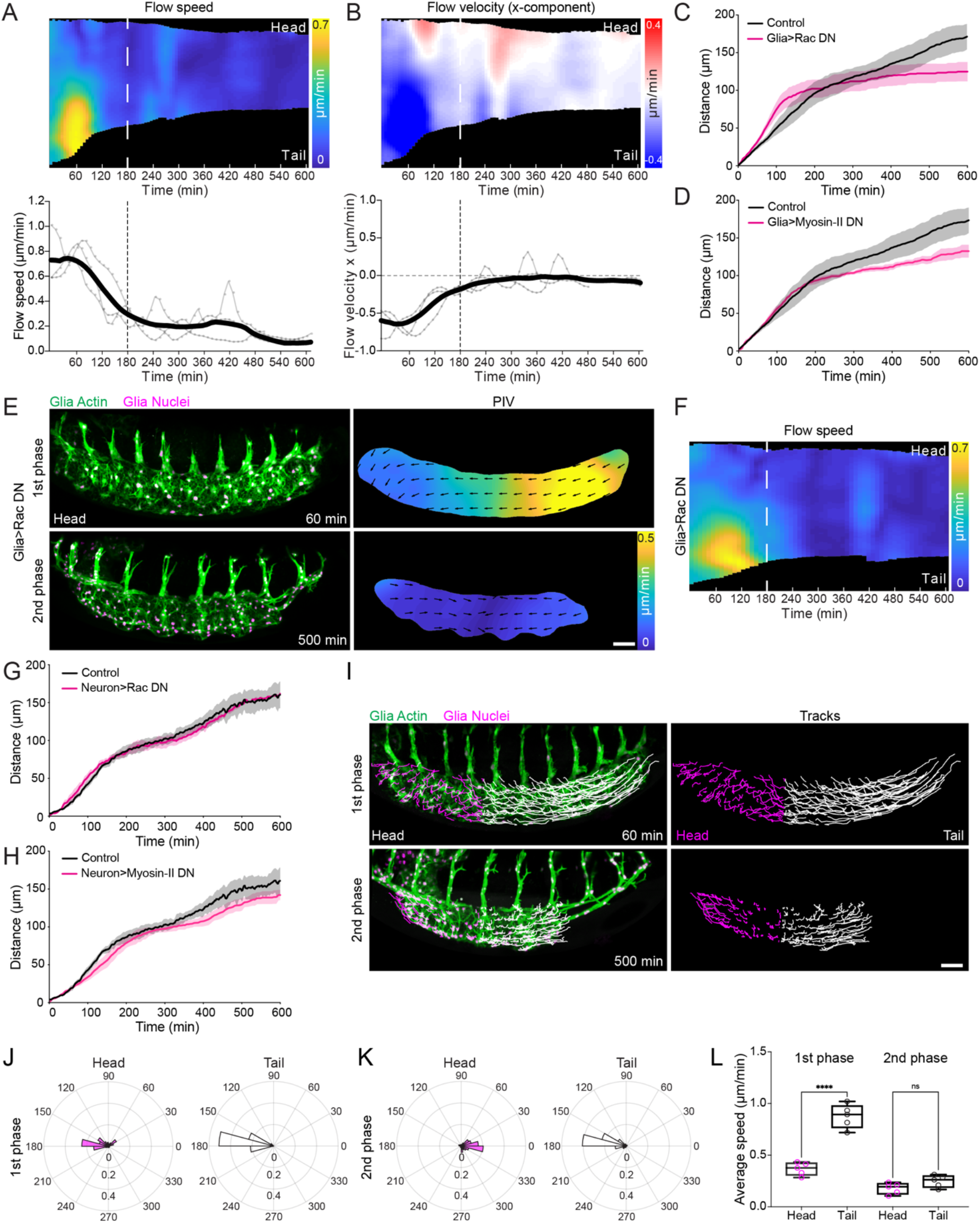
*Drosophila* VNC condensation consists of two distinct temporal stages and initial anisotropic changes in tissue morphology are independent of VNC cellular activity. **(A)** (top panel) Kymograph of the average speed of VNC condensation from head to tail of the tissue from PIV analysis as shown in Fig 1A. Note the asymmetric increase in speed in the tail of the tissue during the first 3 hours of condensation. (bottom panel) Timecourse of the average speed of VNC condensation as measured above highlighting a transition from fast to slow phases around 3 hours. N = 3 embryos. **(B)** (top panel) Kymograph of the average x-component of the velocity along the length of the tissue from head to tail during VNC condensation. Note the predominant negative velocity in the first 3 hours of condensation, which highlights specific motion of the tail of the tissue towards the head. (bottom panel) Timecourse of the average velocity of the x-component as measured above highlighting that during the first 3 hours condensation is predominantly anisotropic from tail to head (*i.e*., the x-component is negative). After 3 hours the average x-component approaches zero suggesting that this longer phase of condensation is symmetric (*i.e*., equivalent from the head and tail of the tissue). N = 3 embryos. **(C,D)** Quantification of VNC condensation by tracking the tail of the tissue as in Fig. 1D,E after expressing dominant negative (DN) Rac (C) or DN Myosin-II (D) in glia reveals that the rate is only affected during the 2^nd^ phase of condensation. Control data is the same in both panels (N = 4 embryos for control, N = 3 embryos for Glia>Rac DN and Glia>Myosin-II DN). **(E)** Live imaging of VNC morphogenesis as in Fig. 1A while expressing Rac DN in glia reveals the presence of an anisotropic 1^st^ phase of condensation. Scale bar = 30 μm. **(F)** Kymograph of the average speed of VNC condensation from head to tail of the tissue from PIV analysis in panel (E). Note the asymmetric increase in speed in the tail of the tissue during the first 3 hours of condensation despite the expression of Rac DN. **(G,H)** Quantification of VNC condensation by tracking the tail of the tissue as in Fig. 1D,E after expressing dominant negative (DN) Rac (G) or DN Myosin-II (H) in neurons reveals little if any effect on VNC condensation (N = 3 embryos for each sample). **(I)** Glial cell motion in the head (magenta) vs tail (white) of the tissue tracked during the 1^st^ and 2^nd^ phases of condensation. Scale bar = 30 μm. **(J,K)** Quantification of the direction of tracks in panel (I) reveals that the cells predominantly move in a tail to head direction (towards 180°) during the first phase (J). In contrast, during the isotropic 2^nd^ phase of condensation the cell tracks are predominantly moving symmetrically towards the center of the VNC (K). N = 3 embryos. **(L)** Quantification of average cell speed during the 1^st^ and 2^nd^ phases of condensation reveals that the motion of cells is fastest within the tail of the tissue during the 1^st^ phase of condensation, while showing no local difference in speed during the 2^nd^ phase. Two-way ANOVA and Tukey’s multiple comparisons test. N = 5 embryos.

**Figure S2.**
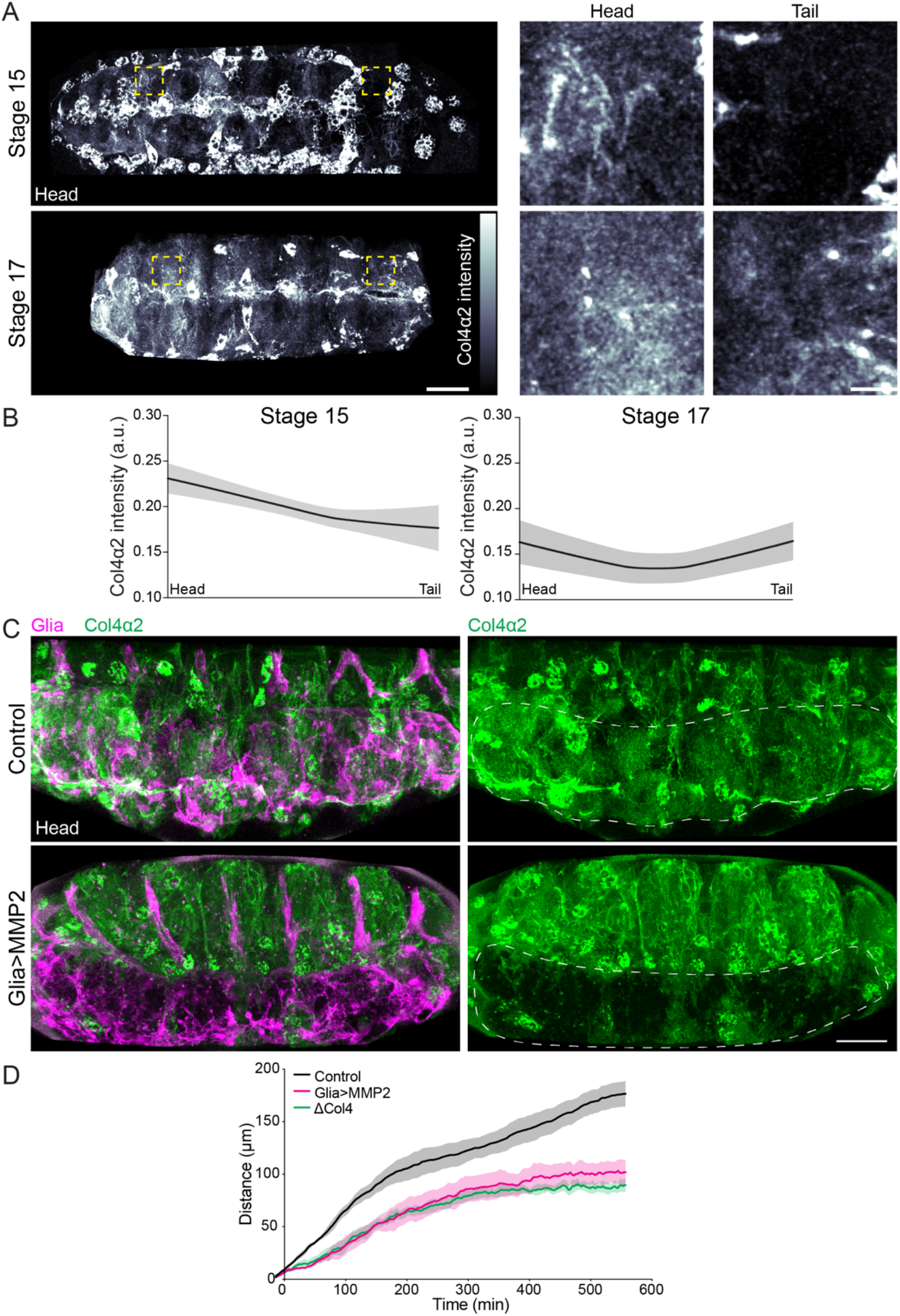
Col4 accumulates along the surface of the VNC and forms a transient gradient during the 1^st^ phase of condensation, and local disruption of the Col4 network around the VNC phenocopies defects observed in Col4 mutants. **(A)** Live imaging of Col4 accumulation on the surface of the VNC during the 1^st^ phase (stage 15) and 2^nd^ phases (stage 17) of condensation. Right panels are high magnification views of the regions highlighted by the yellow squares. Scale bars = 30 μm (left panels) or 5 μm (right panels). **(B)** Quantification of Col4 intensity from head to tail of the VNC reveals a transient gradient during the 1^st^ phase of condensation. N = 3 embryos for each stage. **(C)** Live imaging of Col4 and glia in control embryos and embryos driving MMP2 specifically in glia. Note the local disruption of Col4 accumulation surrounding the VNC (white outline). Scale bar = 30 μm. **(D)** Quantification of VNC condensation by tracking the tail of the tissue as in Fig. 1D,E in *col4* mutants or after expressing MMP2 in glia reveals similar effects on the rate of condensation. Control and *col4* mutant data reused from Fig. 2B (N = 4 embryos for control, N = 3 embryos for Glia>MMP2 and ΔCol4).

**Figure S3.**
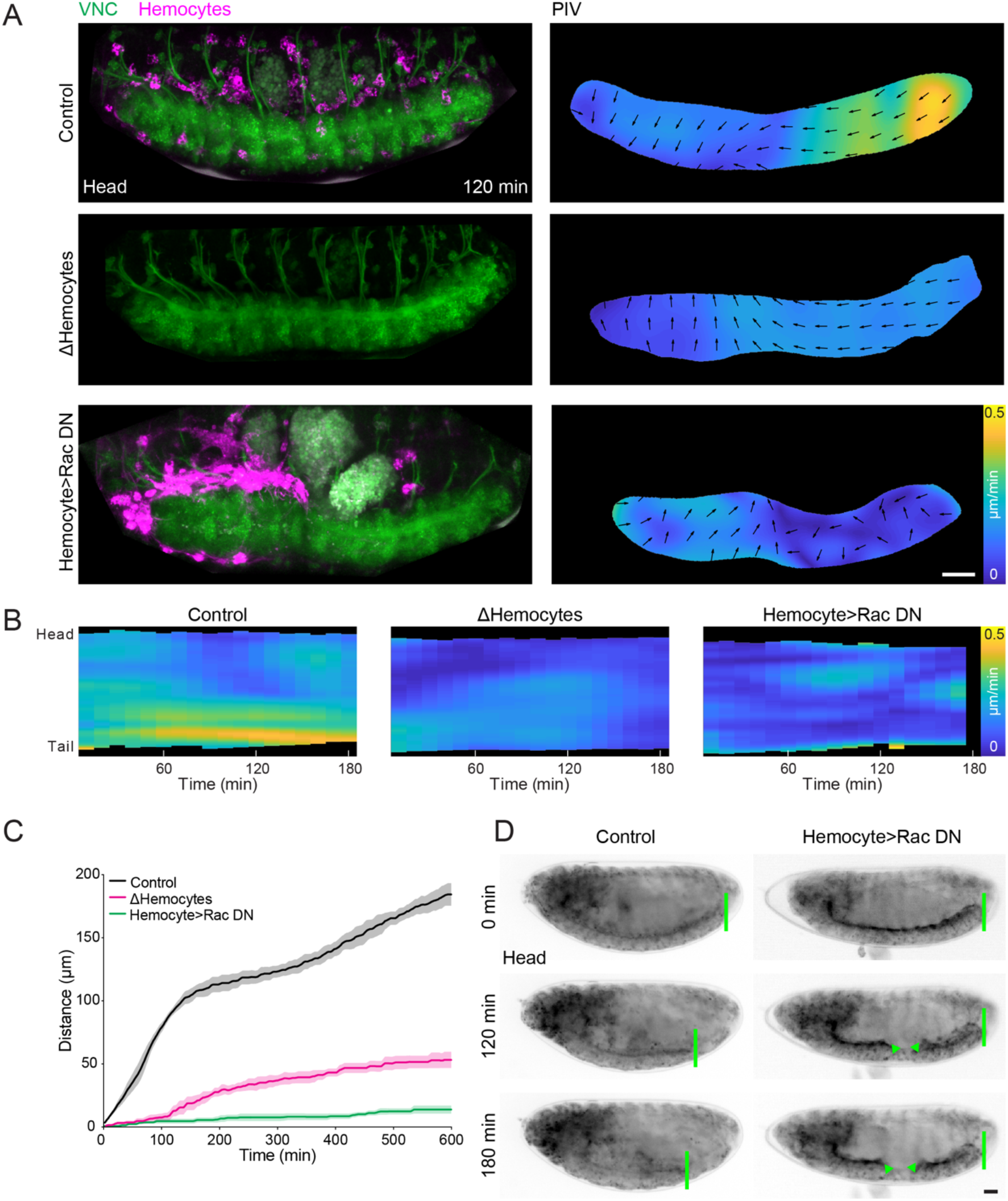
Inhibiting hemocyte migration severely affects VNC condensation. **(A)** Live imaging of VNC morphogenesis (left panels) and PIV (right panels) during the 1^st^ phase of condensation as in Fig. 1A in control, deletion of hemocytes, and expression of Rac DN specifically in hemocytes, which leads to their accumulation in the head of the embryo. Scale bar = 30 μm. **(B)** Kymograph of the average speed of VNC condensation from PIV analysis in (A) highlighting the absence of an anisotropic phase of condensation when perturbing hemocytes. **(C)** Quantification of VNC condensation by tracking the tail of the tissue as in Fig. 1D,E in the genotypes highlighted in panel (A) reveals that loss of hemocytes or inhibiting their migration leads to severe defects in the rate of VNC condensation. N = 3 embryos for each sample. **(D)** Timelapse series of VNC condensation (green lines) in controls and embryos expressing Rac DN specifically in hemocytes. Note that VNC condensation is completely inhibited as the VNC is severely deformed and appears to sever in the center of the embryo (arrowheads). Scale bar = 30 μm.

**Figure S4.**
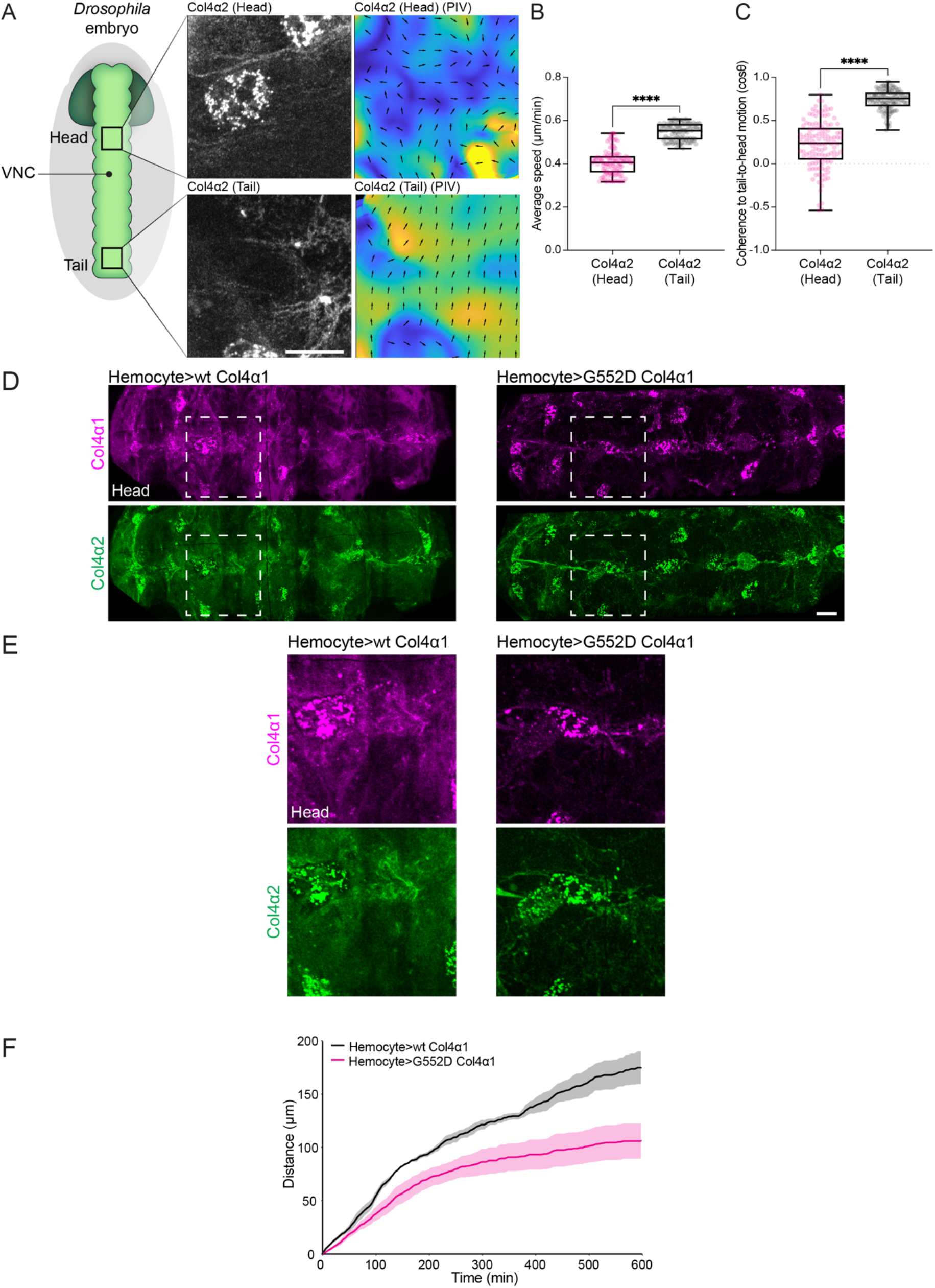
Coherent flow of Col4 along the surface of the VNC is towards the head of the tissue, and driving a G552D temperature sensitive Col4 point mutant transgene specifically in hemocytes severely inhibits VNC condensation. (**A**) (Left panels) Col4 was live imaged during the 1^st^ phase of VNC condensation simultaneously in the head and tail regions of the tissue. (Right panels) Tracking of Col4 motion by PIV in head vs. tail regions of the VNC. Scale bar = 10 μm (**B**) Quantification of the average Col4 speed for each frame reveals increased motion in the tail of the VNC when compared with the head. Mann-Whitney test. N = 127 frames for each sample. (**C**) Correlation of the global alignment and orientation of PIV vectors for each frame reveals that the motion of the Col4 network is more coherent in the tail of the tissue and oriented towards the head (cosθ of 1 represents a tail-to-head orientation). Mann-Whitney test. N = 127 frames for each sample. (**D**) Expression of wild-type or a G552D point mutant Col4α1 in hemocytes during VNC condensation in the background of endogenously GFP-tagged Col4α2. Scale bar = 30 μm. (**E**) High magnification view of the regions highlighted in (D) reveals that the G552D transgene fails to incorporate into the ECM network and also inhibits incorporation of Col4α2. Scale bar = 10 μm. (**F**) Quantification of VNC condensation by tracking the tail of the tissue as in Fig. 1D,E in the genotypes highlighted in panel (D) reveals that expression of the G552D transgene specifically in hemocytes leads to a severe defect in VNC condensation. N = 3 embryos for Hemocyte>wt Col4α1, N = 5 embryos for Hemocyte>G552D Col4α1.

## Supplementary Movie Legends

### Movie 1. VNC condensation involves two distinct phases and the initiation of VNC morphogenesis correlates with hemocyte migration and Col4 deposition, related to Figures 1 and S1

**Part 1. Live imaging of VNC condensation and tracking tissue deformation by PIV highlights distinct phases of condensation.** (Left panel) Time-lapse imaging of VNC condensation in embryos containing glia labeled for actin and nuclei. (Right panel) PIV of VNC condensation reveals that the first ~3hrs of VNC condensation is asymmetric from tail to head. After the first 3hrs condensation is predominantly isotropic as the tissue shrinks symmetrically.

**Part 2. Tracking glial motion during VNC morphogenesis highlights distinct phases of condensation.** Automatic tracking of glia in the head (magenta tracks) and tail (white tracks) of the VNC. Note that during the first ~3hrs of VNC condensation cell motion is rapid and predominantly in a tail to head direction. In contrast, during the remainder of VNC morphogenesis cells are slower with movement towards the center of the tissue.

**Part 3. Live imaging of VNC morphogenesis and Col4 expression by stereomicroscopy reveals condensation coincides with the induction of Col4.** Simultaneous live imaging of VNC condensation and induction of Col4 levels on a stereomicroscope using an endogenous GFP-trap in Col4α2. Example movie was used for quantification of condensation and Col4 levels in Fig. 1D,E.

**Part 4. Live imaging of VNC morphogenesis and Col4 expression by confocal microscopy reveals condensation coincides with the induction of Col4.** Simultaneous live imaging of VNC condensation and induction of Col4 levels by confocal microscopy using an endogenous GFP-trap in Col4α2. Note that as VNC condensation starts, Col4 rapidly begins to assemble on tissues throughout the embryo.

**Part 5. Hemocyte embryonic dispersal correlates with the initiation of VNC condensation.** Simultaneous live imaging of VNC (green) condensation and the embryonic migration of hemocytes (magenta) revealing that condensation coincides with the even hemocyte dispersal within the embryo. Time stamp hh:mm.

### Movie 2. BM mutations and perturbation of hemocytes prevents the initiation of VNC condensation, related to Figures 2 and S3

**Part 1. Live imaging of VNC condensation in *laminin, perlecan*, and *col4* mutants.** (Left panels) Time-lapse imaging of VNC morphogenesis during the 1^st^ phase of condensation in embryos containing glia labeled for actin and nuclei in *laminin, perlecan*, and *col4* mutants. Note the specific disintegration of the VNC in the absence of Laminin. (Right panels) PIV of VNC condensation in the different mutants reveals that *laminin* and *col4* mutants specifically lack an asymmetric 1^st^ phase of condensation.

**Part 2. Live imaging of VNC condensation after perturbation of hemocytes.** (Left panels) Time-lapse imaging of hemocyte migration (magenta) and VNC condensation (green) in control embryos, embryos lacking hemocytes, and embryos containing hemocytes expressing Rac DN to inhibit their migration from the head of the embryo. Note the herniation of the gut specifically during inhibition of hemocyte migration, which appears to severely deform the VNC. (Right panels) PIV of VNC morphogenesis after perturbation of hemocytes reveals a severe defect in the 1^st^ phase of condensation.

**Part 3. Live imaging of VNC condensation after perturbation of hemocyte migration.** Timelapse imaging of VNC condensation in control embryos and embryos containing hemocytes expressing Rac DN to inhibit their migration, which leads to hemocytes residing in the head of the embryo. Note the complete inhibition of VNC condensation after inhibition of hemocyte migration and an apparent severing of the VNC in the middle of the embryo.

### Movie 3. The initiation of VNC condensation correlates with coherent Col4 flow along the tissue surface in a tail to head direction, related to Figure 3 and S4

**Part 1. Live imaging of glial activity and Col4 motion on the VNC surface.** (Top panels) Timelapse imaging of glia containing labeled actin and Col4α2-GFP. (Bottom panels) PIV of glia and Col4 motion. Note the coherent motion of Col4 from tail to head without any noticeable local remodeling by the underlying glial cells.

**Part 2. Live imaging of Col4 motion on the VNC surface in the head vs. tail of the tissue.** (Left panels) Simultaneous time-lapse imaging of Col4α2-GFP on the VNC surface in the head vs. tail of the tissue. (Right panels) PIV of Col4 motion in the head vs. tail of the tissue. Note that in the tail of the VNC there is a predominant movement of Col4 towards the head of the tissue. Movies are oriented with the head at the top of the image. Time stamp hh:mm:ss.

